# Modeling and Cost Benefit Analysis to Guide Deployment of POC Diagnostics for Non-typhoidal *Salmonella* Infections with Antimicrobial Resistance

**DOI:** 10.1101/384933

**Authors:** Carrie Manore, Todd Graham, Alexa Carr, Alicia Feryn, Shailja Jakhar, Harshini Mukundan, Hannah Callender Highlander

## Abstract

Invasive non-typhoidal *Salmonella* (NTS) is among the leading causes of blood stream infections in sub-Saharan Africa and other developing regions, especially among pediatric populations. Invasive NTS can be difficult to treat and have high case-fatality rates, in part due to emergence of strains resistant to broad-spectrum antibiotics. Furthermore, improper treatment contributes to increased antibiotic resistance and death. Point of care (POC) diagnostic tests that rapidly identify invasive NTS infection, and differentiate between resistant and non-resistant strains, may greatly improve patient outcomes and decrease resistance at the community level. Here we present for the first time a model for NTS dynamics in high risk populations that can analyze the potential advantages and disadvantages of four strategies involving POC diagnostic deployment, and the resulting impact on antimicrobial treatment for patients. Our analysis strongly supports the use of POC diagnostics coupled with targeted antibiotic use for patients upon arrival in the clinic for optimal patient and public health outcomes. We show that even the use of imperfect POC diagnostics can significantly reduce total costs and number of deaths, provided that the diagnostic gives results quickly enough that patients are likely to return or stay to receive targeted treatment.

## Introduction

Invasive Salmonellosis can be caused by *Salmonella enterica* serovar Typhi or Paratyphi A and B, S. Paratyphi C, or invasive non-typhoidal Salmonella (NTS) serotypes, including S. Enteriditis and S. Typhimurium. Together, these species are responsible for a gamut of infections from gastroenteritis, typhoid fever, enteric fever to septicemia. NTS, the focal organism of our study, are a major global threat afflicting an estimated 93 million people annually worldwide^1^. The manifestation of NTS infection can vary considerably from mild gastroenteritis to sepsis. Manifestation of NTS infection is divided into invasive and non-invasive disease, of which the former is responsible for approximately 3.4 million illnesses and over 600,000 deaths annually world-wide^2, 3^. Microbiologically treated invasive NTS disease can have a case fatality rate of 20-47% in African adults and children, and accounts for around 39% of community-acquired blood stream infections in sub-Saharan Africa^4–6^. Infants and small children and immuno-compromised individuals such as those with HIV infection or pregnant women, are at highest risk for NTS due to their compromised or naïve immune system. The focus of this study will be on immuno-naïve and immuno-compromised populations that are at high risk for invasive NTS.

A novel genotype of Salmonella enterica subsp. enterica serovar Typhimurium (multi-locus sequence type [ST] 313)^7^ is increasing in prevalence in Sub-Saharan Africa, and different from ST −19 which is predominant in the rest of the world. These strains are associated with predominantly the invasive form of the disease, behave differently from classical NTS strains and are^3^ evolving to transmit between people directly. The outbreaks are also often associated with increased prevalence of HIV^5–7^. Malawi saw an outbreak of NTS from 1998-2004 resulting in 4,956 reported cases of invasive bacteremia disease in Blantyre, a city of about a million people^6^. In a pediatric cohort in Siaya, Kenya, the pediatric prevalence of bacteremia was 11% and 20% of the total pediatric deaths observed in the study were due to bacteremia and 15% of the deaths to *Salmonella* in particular^8^. More recently, ST313 has become increasingly resistant to first-line antibiotics and often exhibits multi-drug resistance (MDR), which is associated with a second wave of outbreaks^6, 7, 9^. Multi-drug resistant NTS represents a threat not only to health in sub-Saharan Africa, but to the world.

Current diagnostics for Salmonella infections, including invasive NTS, are inadequate to guide timely surveillance and decision making. Blood culture, which takes 1-5 days to result and has low sensitivity in clinical samples, remains the gold standard for diagnosis of NTS. Further, culture methodologies require laboratory resources and trained technical personnel, which is not always readily available in resource limited provinces. Immunoassays are available that either target a pathogen-specific antigen (antigen test) or measure the antibody responses to the pathogen (serological tests). Antigen tests have moderate sensitivity and variable specificity, depending on the choice of the recognition ligand (such as antibody) used for the assay. Rapid serological assays are often used for diagnosis of invasive NTS^10^, but these methods are unsuitable for use in regions endemic for NTS, as most individuals from such regions are exposed to the pathogen and will demonstrate an antibody response. For instance, a seroprevalance study in Malawi showed that all infants were exposed to NTS by 16 months of age, and had anti-salmonella IgG antibodies in blood, and would consequently test positive with a serology-based diagnostic irrespective of whether they were infected or not. Polymerase Chain Reaction (PCR) assays offer greater specificity of detection in some cases, but require laboratory resources, cold-chain reagents and trained personnel. They are also known to demonstrate variable sensitivity of outcomes, especially in culture-negative cases. Further, high specificity PCR reactions may be ineffective in rapidly evolving antimicrobial strains of the pathogen.

Better diagnostic tests are needed to improve case finding and management and disease surveillance. Underestimation of NTS prevalence is common due to inadequate and un-affordable diagnostics. There has been a dearth of methods for the effective surveillance, diagnosis and treatment of invasive NTS infections in resource limited areas of the world^11^. Further, Nadjm et al. showed that current WHO guidelines were unable to diagnose almost one-third of children with invasive bacterial disease frequently caused by NTS^12^. Further, half of the isolates were also shown to demonstrate antimicrobial resistance, requiring further characterization to warrant effective treatment. It is important to note that given the acute manifestation of NTS infections, including invasive disease, any diagnostic test performed to guide intervention should be rapid and usable at the point of care (POC). However, POC diagnostics for use in resource poor regions of the world such as sub-Saharan Africa, where the disease is endemic, should be able to operate with minimal power requirements, technical expertise, and laboratory infrastructure, while being inexpensive and robust for use. Triage diagnostics, even with lower sensitivity but a rapid time to result, can facilitate decision making and treatment at the POC. Indeed, the return to the clinic for securing diagnostics information and initiating treatment is not an option for many individuals in resource poor regions^13^. The absence of such specific rapid diagnostics to identify causal organisms and further characterize antimicrobial strains is one of the major limitations to effective treatment of invasive NTS, thereby minimizing associated mortality and morbidity. Indeed, such rapid POC diagnostic tests have already proved to be cost effective in regions endemic for malaria and with high multidrug resistance^9^. Therefore, the intent of this study is to determine when it is most beneficial to deploy a diagnostic for NTS infection, which type of diagnostic test is best to guide situational awareness and improve patient outcome, and outline these findings in a cost benefit analysis.

A major reason for timely and targeted treatment is to prevent mortality associated with antibiotic resistance, and also minimize its further spread by use of inappropriate treatment strategies. For the purposes of this discussion, antibiotic resistance is defined as resistance of a bacteria to a broad spectrum antibiotic that was originally effective for treatment of the disease in question (aka NTS). The improper use of antibiotics can accelerate emergence of resistant bacterial strains. In a study in Ghana, rates of NTS resistance to particular antimicrobials ranged from 25% to 62%, with 54% of strains being multi-resistant^14^. In Congo, 80% of NTS strains sampled were multi-drug resistant^15^. In Kenya, where this study is based, NTS antibiotic resistance rates were found to be similar, with more than half of the isolates demonstrating resistance to at least one common antibiotic, and 74% demonstrating multi-drug resistance^16^. The choice of antibiotics and the treatment regiment used are completely different for drug-sensitive, resistant and multi-drug resistant manifestations of the disease. Broad spectrum antibiotics commonly used to treat NTS such as chloramphenicol, ampicillin, gentamicin, ciprofloxacin, and trimethroprim-sulfamethoxazole^6, 16^ are replaced by more expensive and elusive third generation cephalosporins and fluoroquinolones^5^ in drug-resistant manifestations of the disease. Without correct treatment, resistance proliferates and death rates increase. Effective diagnostics at the POC, evaluation of antimicrobial resistance, and monitoring prognosis can facilitate targeted treatment of the disease, saving lives and minimizing further spread of dangerous resistant phenotypes within the community.

While diagnostics can be helpful in determining the correct therapeutic strategy for a given patient, minimizing the misuse of antibiotics, and thereby controlling the emergence of drug resistance, it is unclear whether the added expense, time, and questionable dependability of some diagnostic methods out-weigh these benefits. The purpose of our work is use mathematical modeling to determine the answer to this question. Such models have been developed, albeit with variable focus and for other pathogens, before. Mathematical models of typhoid are reviewed in^17, 18^, including a landmark model by^19^, highlighting the need for models and economic analysis. Feasey et al. considered the interaction between malaria, HIV, malnutrition, and rainfall, and pediatric invasive NTS (iNTS)^20^. They used a structural equation model to observe the effect that each risk factor made in the decline of pediatric iNTS with response to public health investments and the connection amongst the risk factors. In the current literature, there are very few mathematical models for *Salmonella*, and none for NTS at a population scale. Models that explicitly include point of care (POC) diagnostics and their impact on patient and community outcomes are also uncommon. Thus, the goal of our work is to develop a comprehensive mathematical approach to derive cost-benefit analysis for the development and deployment of diagnostics, considering all parameters that influence outcome (acuteness of infection, economic stability of the population in consideration, nature of the population, endemicity of the pathogens, immuno-compromised versus healthy population and others) for invasive NTS infections.

To develop and evaluate the model, we identified the most common diagnostic approaches (not all-inclusive, but predominant) for NTS infection. We derived the associated costs for these selected diagnostic approaches considered from literature, as described here. Neither the list of diagnostics nor the costs are intended to be exhaustive, but are used to provide a relative, but accurate, estimate of cost versus benefit, when employed in the model- thus demonstrating a methodology for making such assessments quantitatively and accurately. Diagnostics considered in this study include Polymerase Chain Reaction (PCR), bacterial culture (BC), and antibody-based tests (primarily serological measurements). PCR tests cost about 10 USD, require 24 hours to return results, and are approximately 90 percent accurate^10, 21^. Bacterial cultures cost about 5 USD, also require nearly 24 hours to results, and are less reliable than PCR, (40-80 percent accuracy rate, depending on the situation)^10, 22, 23^, and even lower (∼30 percent) for fecal samples^10, 22^. Fecal culture cannot discriminate gastro-intestinal manifestation of the disease from invasive Salmonellosis. Antibody tests cost about 1 USD, require 15 minutes to results, and range in accuracy from 78 to 100 percent^10, 22, 24^. As such, antibody based tests which largely satisfy the requirements of POC use with respect to simplicity and cost are lacking in specificity, and are unable to discriminate between invasive and gastrointestinal disease, and are associated with high false-positive rates in endemic populations. In fact, a seroprevalence study of healthy children in Malawi revealed that they all had anti-Salmonella IgG antibodies by the age of 16 months, which suggests that infants have been universally exposed to either non-typhoidal salmonellae or cross-reactive antigens at a young age^25^. Of course, specific costs, sensitivity and specificity rates, and time to return results will vary across companies providing the tests and the tests themselves, so we used the prices and turn-around times listed here as generic estimators of relative costs. Given these expenses, the intent of this study is to consider when it is most beneficial to deploy a differential diagnostic, which diagnostic is best, and outline these findings in a cost-benefit analysis. Such analysis can be used in future when new diagnostic methods are being considered for development and deployment to assess whether the technology can truly impact care in a given situation.

In many regions of the world, antibiotics are prescribed according to symptom severity rather than according to the results of an empirical diagnostic^22^. Our model and analysis is based on the at-risk population size and protocols of the clinic systems in Siaya, Kenya^8^. This area is representative of regions with poor infrastructure and health access as well as high at-risk populations in which NTS seems to thrive. Health authorities in Kenya suggest no antibiotic be given to patients with gastrointestinal symptoms unless a fever is also present, rather administering supportive care such as oral re-hydration salts and fluids to mildly symptomatic patients^16, 26, 27^. Antibiotic treatment is not usually advised for uncomplicated cases because there is no evidence that the recovery rate is accelerated, therefore the patient outcome would be similar^28^. Also, unnecessary use of antibiotics can cause negative side effects in the individual, and promote antimicrobial resistance development^6^. However, some studies have shown that an individual can be a potential carrier for Salmonella infection for a longer period of time without an extended treatment regimen^22^, while others suggest prolonging carrier shedding with treatment^29^. Antibiotic treatment may be necessary for infants, elderly and immuno-compromised patients even if they are mildly symptomatic^29^. Patients with fever are often given broad-spectrum antibiotics (unless misdiagnosed as malaria, and antimalarials are initially prescribed). A severely symptomatic patient needs to be treated properly and quickly because enteric fever has a high mortality rate, upwards of 30%, without proper treatment. The mortality rate drops to as low 0.5% when the correct treatment is given^5^. Thus it is imperative to treat patients with invasive fever quickly. With the rapid increase in multi-drug resistance, and the prevalence of HIV co-infection, tailored and targeted treatment is critical for survival, and for minimizing the long term impact on the population. We assume in this study that when patients arrive to the clinic they are classified as either “mildly symptomatic” or “severely symptomatic”, where “mildly symptomatic” patients experience diarrhea and gastrointestinal symptoms only, and “severely symptomatic” patients experience fever, suggesting invasive infection.

In many developing nations, people are able to obtain and administer antibiotics without prescriptions from a medical professional^16, 30, 31^. For example, in Kenya, 24% of children under 5 were reported to have fever in the past two weeks, 55% of them sought medical care, and 44% received antibiotics. Similarly, 16% of children under 5 were reported to have diarrhea in the past 2 weeks, 56% sought medical care, and 17% received antibiotics^31^. While this current approach is the most immediately accessible and initially inexpensive, it allows antibiotics to be administered haphazardly without regard to the antibiotic’s ability to treat the present strain. Such improper administration of antibiotics results in increased antibiotic resistance. Accordingly, different combinations of antibiotic treatment and diagnostic deployment may be better able to achieve the goal of efficiently minimizing the infectious population and improving patient outcomes.

Herein, we report the development of our model for NTS dynamics at a population level, and then demonstrate application of the same to various relevant scenarios. We used an Susceptible-Infectious-Recovered (SIR) type mechanistic population-level model to allow us to assess how use of diagnostics affects disease dynamics, total infection rate, and progression, and deaths caused by NTS in at-risk populations. A cost-benefit analysis was applied to determine which scenarios limited the overall costs. To our knowledge, this is the first population-level mathematical model of NTS and the first mathematical model that incorporates diagnostics and targeted treatment for NTS.

### Scenario Setup

The primary goal of this modeling effort is to determine the costs and benefits of deploying POC diagnostics for the control of NTS while minimizing antimicrobial resistance in high-risk groups. The potential for POC diagnostics to improve patient care is considered through a series of scenarios differentiated by which group of infectious individuals receive a diagnostic. Infectious individuals are classified by the severity of their symptoms to form the different groups: **mildly symptomatic** (gastroenteritis) and **severely symptomatic** (invasive disease with fever). As stated before, these scenarios are meant to illustrate the utility of the model under multiple scenarios and provide motivation for investment in POC diagnostics, thus are not exhaustive. The distinction between symptom intensity determines which patients receive diagnostics in each of the four scenarios:

1. *Full Diagnostic Deployment*. All patients receive a POC diagnostic, and are prescribed targeted treatment based on diagnostic results. They are given alternative treatment in the event of false positives or false negatives exhibited by failure to recover.
2. *Partial Deployment of Diagnostics*. POC diagnostics are only given to mildly symptomatic patients with gastroenteritis and no fever, and targeted antibiotics are prescribed as determined by the diagnostic results. Severely symptomatic patients with fever are *immediately* empirically prescribed the broad spectrum antibiotic treatment without a diagnostic due to often urgent needs of the patient. Then, if they do not respond to that, they are redirected (without a diagnostic) to antibiotics appropriate for the resistant strain.
3. *No Deployment, Antibiotics For All*. Neither mildly symptomatic nor severely symptomatic patients receive a diagnostic, but both groups immediately receive broad-spectrum antibiotic treatment and are redirected to the alternative treatment for the resistant strain if they do not respond to the initial treatment. This assumes that, in contrast to Kenyan guidelines, antibiotics are prescribed for diarrhea or people are self-treating with antibiotics.
4. *No Deployment, Antibiotics For Severely Symptomatic*. Neither mildly symptomatic nor severely symptomatic patients receive a diagnostic. Severely symptomatic (invasive) patients receive a broad spectrum antibiotic and patients that do not recover are given a treatment appropriate for the resistant strain. Mildly symptomatic with gastroenteritis only are not prescribed antibiotics and do not self-treat.

The first scenario assumes that a POC diagnostic is available and regularly used for both mildly symptomatic and severely symptomatic individuals. The second scenario assumes that a physician will not wait for diagnostics to begin antibiotic treatment for the very ill severely symptomatic individuals, but may order diagnostics for those who are ill but not yet serious enough for hospitalization. The diagnostics require an added cost and initial expense and likely result in more prescriptions of the costly resistant treatment as opposed to the scenarios where the less expensive broad spectrum antibiotic is given without heed to resistance. However, if it can be shown that use of a diagnostic efficiently minimizes the infectious population and deaths resulting from NTS disease, this would contribute to lessened long-term expenses and provide motivation for diagnostics to be strategically deployed. The third and fourth scenarios are most common in Kenya^16, 26^. While physicians are recommended not to treat mildly symptomatic cases with antibiotics, people can and often do obtain antibiotics without a prescription and some physicians may prescribe antibiotics even though they are not recommended^30, 32, 33^. This would align with the third scenario, while strict adherence to recommendations would correspond more to the fourth scenario. Both assume that diagnostics are either not readily available or are rarely used or not appropriate. The third and fourth do the least to address the concern of antibiotic resistance because of the higher risk of improper treatment.

## Results

We focused the model on NTS spread in immuno-suppressed (immuno-naive and immuno-compromised) individuals in the population. However, we also considered a version of the model that accounted for transmission from otherwise healthy adults and environmental transmission from a water or food source (henceforth referred to as the “Environmental Compartment”, see Supplemental Material for full model equations and analysis). While most healthy adults will be functionally immune to the circulating strains, some may still be susceptible and/or may shed the pathogen. However, available evidence suggests that a very small fraction of healthy adults become infected with and shed NTS. Im et al. 2016^34^ found that prevalence in stool of healthy adults ranged from 6.1 - 17.2 (10.3) per 100k in Senegal and 16.5 - 35.1 (24.1) per 100k in Guinea-Bissau. Feasy et al. 2012^5^ found that only 5% of iNTS cases were in healthy adults, while the rest were HIV-positive. Similarly, Gordon et al 2002^35^ found that 77 of 78 adults with iNTS were HIV positive. Another study estimated that somewhere between 0 - 3.6% of people infected with NTS in developing countries in Africa end up being carriers for several weeks or months^36^. While minimal, transmission from healthy adults (and potentially animals) could have an effect on our analysis. There is very little known about the exact role that environmental transmission plays in NTS in developing nations in Africa that we are considering here (see, e.g. Kariuki et al 2015^37^). While NTS have been found in environmental samples and/or in animals in some regions, the strains found in soil or animals are often not the same as those circulating in the humans^38, 39^. In fact there is evidence that transmission is becoming functionally human-to-human in these low-resource areas^37^. Since animals and humans live in close proximity and often share the same water sources in low-resource areas, we used one compartment to account for all additional sources of infection not coming from the high-risk group. We allowed for a small constant input of infectious doses of NTS to this Environmental compartment to account for long-term shedders. Otherwise, NTS is assumed to pass back and forth between the Environmental compartment and the high-risk group. We found that while the extended model with healthy adults and environmental transmission changed the magnitude of simulated outbreaks, it did not change the qualitative patterns of our results and the relative costs and benefits of deployment of a POC diagnostic. Since general trends of the high-risk only model were preserved by the more complex model (see Supplemental Material Figures 2 and 3), we present results here for the simplified model. This is to preserve interpretability, to minimize uncertainty in parameters related to the Environmental compartment, and because evidence points to most transmission in low-resource countries being effectively direct and primarily in the high-risk population.

We ran simulations for Scenarios 1–4 for each of “antibody”, bacteria culture (BC), and PCR POC diagnostics explicitly considering both drug-sensitive and drug-resistant strains circulating (see Figures 1–2). Our model only considers here cost of point of care (POC) diagnostics that can be used by already available staff and with minimal additional space or resources. Table 1 gives the cost of each possible ordered combination of Antibiotic treatment to which a sensitive strain of the pathogen responds (A), Resistant treatment to which the resistant strain responds (R), and Diagnostic deployment (D) in each of the scenarios. For example, the column heading “Cost DAR” gives the cost of first using a Diagnostic, followed by Antibiotic treatment, followed by the Resistant treatment (this would be an instance of improper treatment). Table 2 gives the number of deaths from NTS (D) after 1,000 days of simulation, in addition to the percentage of improperly treated patients and the total number of diagnostics used.

**Figure 1.**
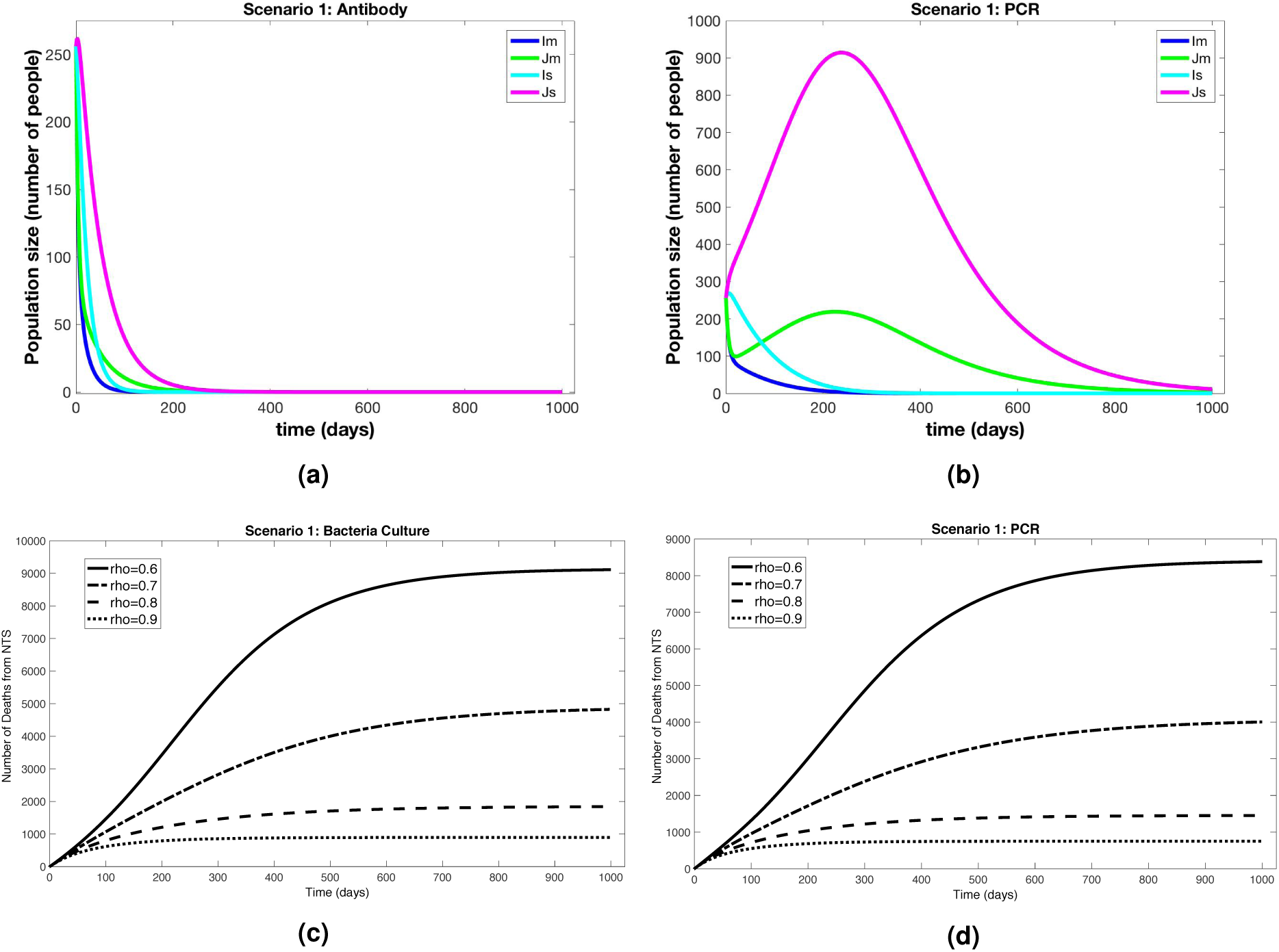
Simulations of Scenario 1 (*Full Diagnostic Deployment*). Figures 1a–1b show outputs from infected compartments of the differential equations for the antibody and PCR diagnostics, respectively. *I*_*s*_ is the resistant strain and severely symptomatic, *J*_*s*_ is the sensitive strain and severely symptomatic, *I*_*m*_ is the resistant strain and mildly symptomatic, *J*_*m*_ is the sensitive strain and mildly symptomatic. Figures 1c–1d show total number of deaths through time for different values of *ρ*, the proportion of patients who stay for diagnostic result and receive targeted treatment, for bacteria culture and PCR.

**Figure 2.**
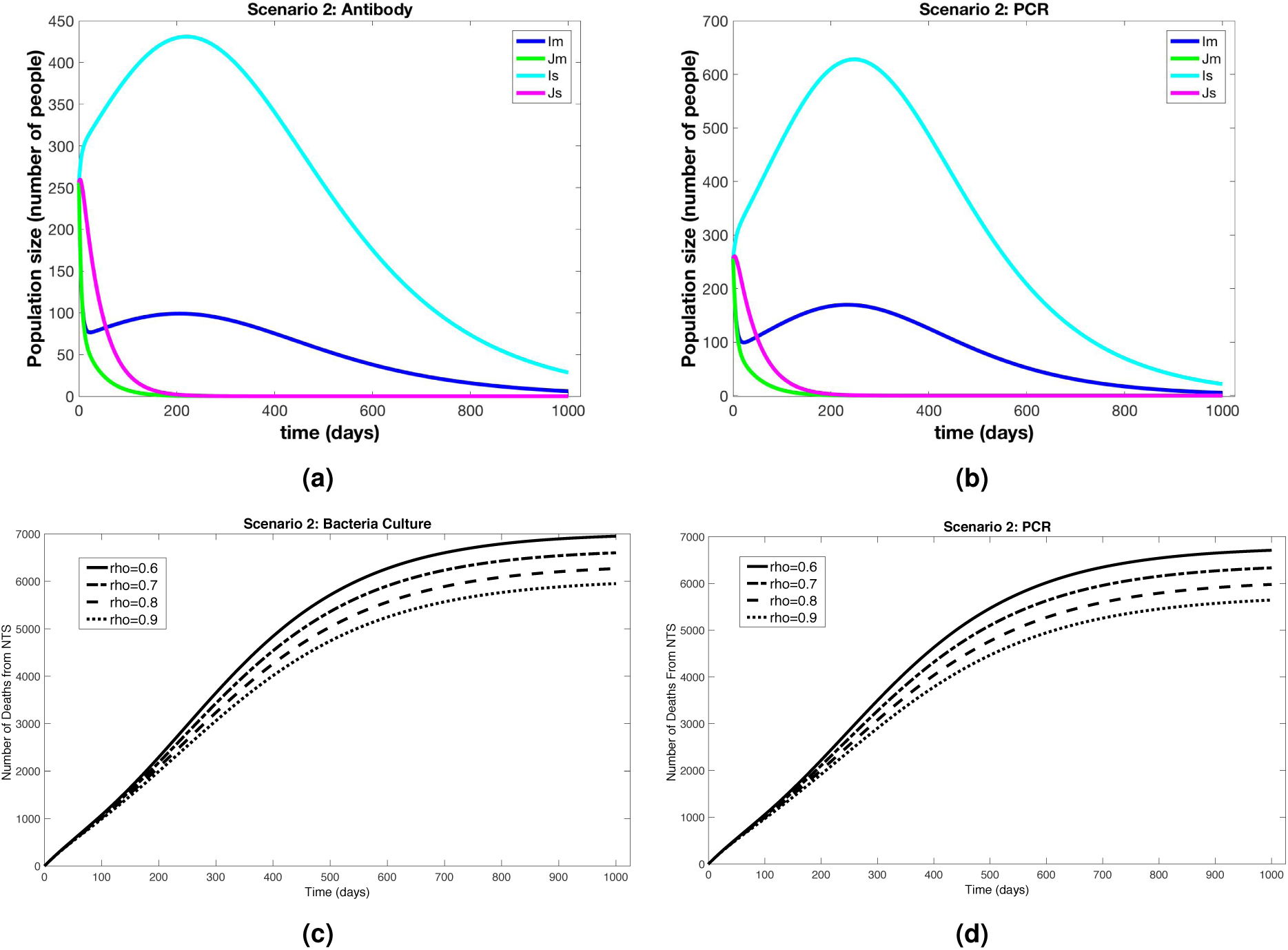
Results from Scenario 2 (*Partial Deployment of Diagnostics*). Figures 2a–2b show outputs from the infected compartments of the differential equations for the antibody and PCR diagnostics, respectively. *I*_*s*_ is the resistant strain and severely symptomatic, *J*_*s*_ is the sensitive strain and severely symptomatic, *I*_*m*_ is the resistant strain and mildly symptomatic, and *J*_*m*_ is the sensitive strain and mildly symptomatic. Figures 2c–2d show total number of deaths through time for different values of *ρ*, the proportion of patients who return for diagnostic results and receive targeted treatment, for bacteria culture and PCR.

**Table 1.** Costs of diagnostic deployment and antibiotic use for each scenario where A is the cost of standard antibiotic treatment (effective on sensitive strain), R the cost of resistant strain treatment, and D the cost of the diagnostics. All costs are in U.S. Dollars (USD).

| Scenario | Diagnostic Used | Cost A (USD) | Cost AR (USD) | Cost DA (USD) | Cost DAR (USD) | Cost DR (USD) | Total Cost (USD) |
| --- | --- | --- | --- | --- | --- | --- | --- |
| 1 | Antibody | 0 | 0 | 12,028 | 6,379 | 47,411 | 65,818 |
| | BC ($\rho = 0.6$) | 0 | 0 | 240,548 | 103,573 | 142,916 | 487,037 |
| | PCR ($\rho = 0.6$) | 0 | 0 | 333,506 | 42,741 | 123,966 | 500,213 |
| | BC ($\rho = 0.8$) | 0 | 0 | 60,565 | 30,657 | 75,981 | 167,203 |
| | PCR ($\rho = 0.8$) | 0 | 0 | 70,963 | 10,169 | 74,143 | 155,274 |
| 2 | Antibody | 7,478 | 534,847 | 2,581 | 12,454 | 309,942 | 867,302 |
| | BC ($\rho = 0.6$) | 8,332 | 723,544 | 2,531 | 17,238 | 257,184 | 1,008,829 |
| | PCR ($\rho = 0.6$) | 8,302 | 698,158 | 3,679 | 7,501 | 313,148 | 1,030,788 |
| | BC ($\rho = 0.8$) | 7,931 | 653,079 | 3,039 | 19,539 | 289,468 | 973,056 |
| | PCR ($\rho = 0.8$) | 7,896 | 622,510 | 4,408 | 8,335 | 346,210 | 989,359 |
| 3 | None | 9,744 | 1,014,473 | 0 | 0 | 0 | 1,024,217 |
| 4 | None | 10,151 | 973,402 | 0 | 0 | 0 | 983,553 |

**Table 2.** Number of deaths from NTS, percent of cases improperly treated, and the number of diagnostics used in each scenario run for 1,000 days.

| Scenario | Diagnostic Used | Number Deaths (People) | Improperly Treated (Percent) | Num. Diagnostics Deployed |
| --- | --- | --- | --- | --- |
| 1 | Antibody | 429 | 4.2% | 2,118 |
| | BC ($\rho = 0.6$) | 9,113 | 6.4% | 21,460 |
| | PCR ($\rho = 0.6$) | 8,382 | 2.6% | 20,383 |
| | BC ($\rho = 0.8$) | 1,840 | 6.7% | 6,056 |
| | PCR ($\rho = 0.8$) | 1,452 | 2.5% | 5,009 |
| 2 | Antibody | 5,141 | 59.1% | 5,329 |
| | BC ($\rho = 0.6$) | 6,952 | 70.3% | 4,226 |
| | PCR ($\rho = 0.6$) | 6,710 | 67.1% | 4,612 |
| | BC ($\rho = 0.8$) | 6,268 | 66.2% | 4,773 |
| | PCR ($\rho = 0.8$) | 5,979 | 62.8% | 5,118 |
| 3 | None | 7,512 | 93.3% | None |
| 4 | None | 9,441 | 92.8% | None |

We found that the status quo baseline scenarios (Scenarios 3 and 4) generally resulted in the highest number of deaths, larger outbreaks, and the highest costs. Because of the prolific but un-targeted use broad spectrum antibiotics in the baseline scenarios, the resistant strains have a distinct advantage over the non-resistant strains as evidenced by the initial increase in number of infected for resistant strain curves (*I*_*s*_ and *I*_*m*_ in Figure 3) while the non-resistant strain dies out relatively quickly. This indicates that, under our model assumptions and parameter values, if resistance were not present, NTS outbreaks would not be sustainable in the human population without long-term zoonotic or human carriers. However, with antibiotic resistance, sustained outbreaks can occur in the human population without outside reservoirs.

**Figure 3.**
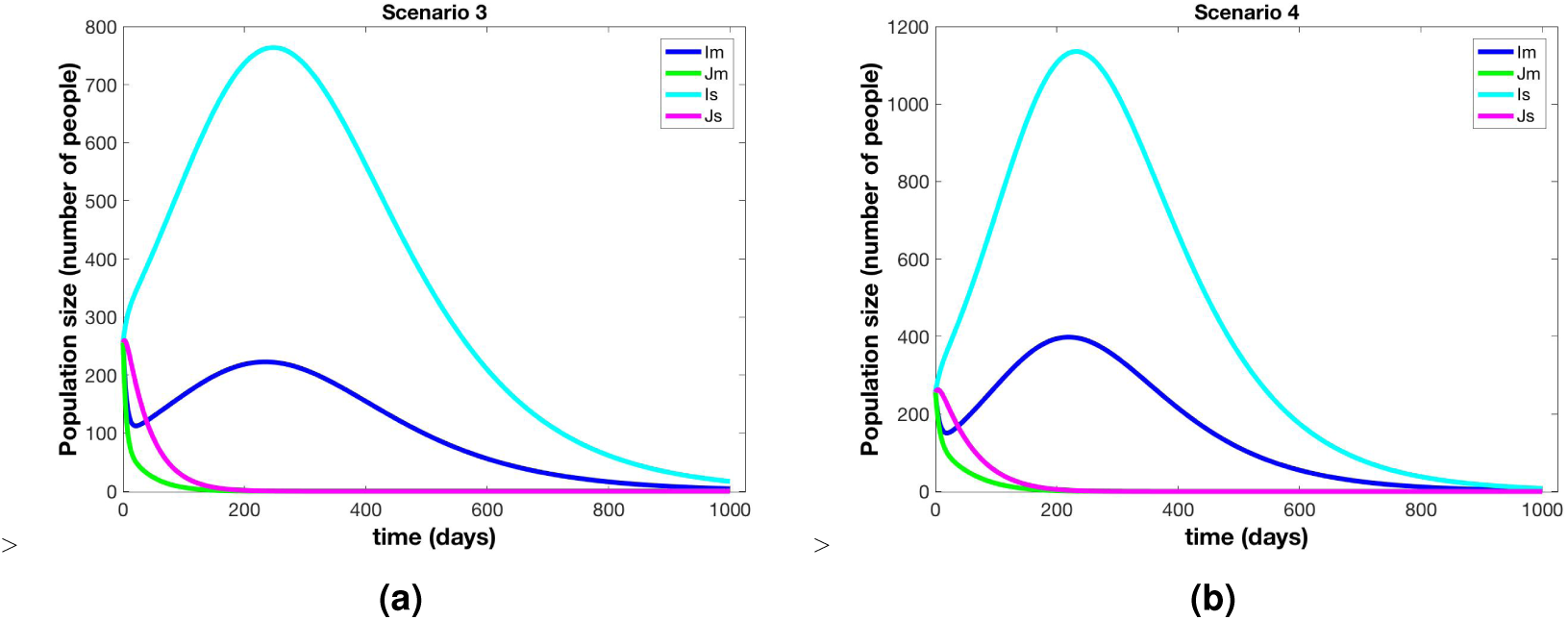
Results from Scenario 3 (*No Deployment, Antibiotics For All*) and Scenario 4 (*No Deployment, Antibiotics For Severely Symptomatic Only*). Figure 3a and Figure 3b do not display the Susceptible or Death from Disease populations in order to better observe the Infectious compartments. *I*_*s*_ is the resistant strain and severely symptomatic, *J*_*s*_ is the sensitive strain and severely symptomatic, *I*_*m*_ is the resistant strain and mildly symptomatic, *J*_*m*_ is the sensitive strain and mildly symptomatic. The outbreak in Scenario 3 is larger than those of Scenario 2 in Figure 2. Scenario 4 results in the largest outbreak compared to all other scenarios.

When diagnostics are partially deployed to mildly symptomatic individuals, the outbreak size is decreased along with a decrease in number of deaths from NTS compared to no diagnostic use (Scenario 2, Figure 2 and Tables 1–2). Costs are also decreased when antibody diagnostics are used, but for culture and PCR, total costs are larger than in Scenario 4 while still less than Scenario 3. So, while partial diagnostic deployment does save lives and decrease NTS outbreak size, it does so at an increased or only slightly decreased total cost (Tables 1–2). The cost per life saved ranges from 200-400 USD for Scenario 2. Importantly, the number of lives saved and costs of POC diagnostics and treatment depend on the parameter *ρ*, which is the proportion of patients that return for diagnostic results and targeted treatment. Since the antibody diagnostic gives results within minutes, we assume *ρ* = 1, or all patients stay. However, since results for BC and PCR may take several hours or a day, we showed results two scenarios, *ρ* = 0.6 and *ρ* = 0.8 in Table 2. When *ρ* = 0.8, the improvement of Scenario 2 over Scenarios 3 and 4 in terms of both cost and number of deaths from NTS is marked. However, gains are moderate but noticeable when *ρ* = 0.6.

When diagnostics are fully deployed (Scenario 1) and directly inform choice of treatment, the outbreak peak is decreased by a factor of 4 compared to Scenario 4 for the antibody diagnostic. For the antibody diagnostic scenario, the outbreak is quenched within the first three months (Figure 1). In scenarios where more time-consuming diagnostics such as PCR and culture were used, results depended on the value of *ρ* (Tables 1–2). Since diagnostics are applied universally in this scenario, it is critical that most patients return or stay to get the diagnostic result and treatment. When *ρ* = 0.8, so 80% of patients return for treatment and results the next day, Scenario 1 is hands-down the best option (Figure 1a). The outbreak is extinguished quickly and costs per life saved range from 7-22 USD. However, if *ρ* = 0.6, then Scenario 2 out-performs Scenario 1 in terms of lives saved (Figures 1c-1d). The total costs, including diagnostics and antibiotics, were significantly less for Scenario 1 than in any other scenario when *ρ* = 0.8. In Scenario 1, when compliance is decreased to *ρ* = 0.6 by having to wait a day for diagnostic results (Figures 1c-1d), 40% of severely symptomatic cases remain untreated, so the number of deaths is comparable to Scenarios 3 and 4.

While data capturing a full outbreak of NTS is rare, we compared our results to a recorded outbreak in a location in Malawi about the same population size as we consider here. To simulate this outbreak of a new strain, we ran scenario 3 with lower initial conditions (5 people in each infectious category). The Blantyre district in Malawi serves a population of about 1 million urban and rural-dwelling people^6^ and the study was over 7 years in the district’s government funded hospital. They recorded a total of 4, 956 cases of invasive NTS during the outbreak over 7 years. The hospital does not treat all sick people in the district, and many who had NTS during this outbreak either did not seek treatment at this hospital, or were not properly diagnosed. While we couldn’t find a health care usage study for Malawi, only 20-30% of the population in Tanzania sought health care at a hospital for fever, with percents throughout Africa ranging from 20-80%^40^. We also know that in high-risk groups about half of NTS cases are invasive (Table 4). Then, we conservatively estimated that the proportion of total NTS cases in the Blantyre high-risk group observed by the hospital is: proportion invasive × proportion seeking care × proportion of population the hospital serves = 0.5·0.3·0.4 = 0.06 or 6%. Then there were likely more than 39, 000 cases of NTS in Blantyre during the outbreak and, with a 25% death rate for iNTS cases, more than 4, 800 deaths. Our model estimated 5, 700 deaths during the seven year outbreak (see Figure 4 in Supplemental Material).

The basic reproduction number, *ℛ*_0_, is the expected number of secondary cases resulting from a single infectious case introduced into a fully susceptible population. It is a measure of the capacity of a pathogen to spread within a population and cause an epidemic. If *ℛ*_0_ < 1 then it is unlikely for the disease to cause an epidemic, while if *ℛ*_0_ > 1, and outbreak is likely. For Scenario 4 (simplified model) with no POC diagnostics used, 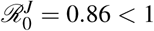 while 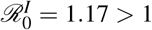. So, in the absence of diagnostics, the sensitive strain will die out while the resistant strain will persist due to large-scale improper use of antibiotics. However, for Scenario 1, with full use of POC diagnostics, and the antibody diagnostic, 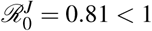 and 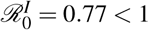, and both strains will die out. We found that the basic reproduction number for each scenario is highly sensitive to the transmission rate, *α*, in a way that depends upon the scenario and diagnostic used (Figure 1 in Supplemental Material). *ℛ*_0_ is also sensitive to the proportion of cases that become invasive, *γ* and *β* for sensitive and resistant strains respectively, with *ℛ*_0_ increasing as both *γ* and *β* increase. Finally, *ℛ*_0_ is sensitive to the compliance rate, *ρ*, which depends on the time it takes to get the diagnostic result back (Figure 1 in Supplemental Material). Understanding how *ℛ*_0_ changes as the parameters change informs both potential avenues of intervention and changes in disease dynamics under particular scenarios.

We also used statistical sensitivity measures (Partial Rank Correlation Coefficients (PRCC) and extended Fourier Amplitude Sensitivity Testing (eFAST)) to understand how the total number of deaths from NTS is related to the parameter values. Since PRCC and eFAST reveal different aspects of how parameters influence a model, for a more complete sensitivity analysis it is good practice to use both sensitivity measures^41^. PRCC values give information as to what extent changing the value of one parameter will increase or decrease the value of the output parameter (where uncertainties in other parameters are discounted). eFAST, on the other hand, provides insight into which uncertainties in parameters cause the most uncertainty in the model output. Thus, the most important set of parameters from PRCC analysis reveal the parameters we should target if we wish to most effectively reduce deaths (*D*), whereas the parameters tagged as important in the eFAST analysis tell us for which parameters we should obtain more accurate values in order to reduce our uncertainty in the output, *D*.

The model output of interest, total number of deaths from NTS, is most sensitive to parameters with large PRCC and/or eFAST sensitivity indices. The parameter *α* (the transmission rate) returns the highest PRCC and eFAST sensitivity indices for all four scenarios. Furthermore, in all scenarios, PRCC and eFAST are in strong agreement regarding the importance of each parameter. In Scenario 1, PRCC and eFAST both rank *κ*_*a*_ (rate of clearance of sensitive strain after receiving antibiotics) as the second most important, followed by *γ* (proportion of non-resistant infections that are invasive). In Scenario 2, PRCC and eFAST return *β* (proportion resistant infections that are severely symptomatic/invasive) and 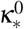 (rate of clearance of the resistant strain after receiving broad spectrum and then resistant-appropriate antibiotics, with no diagnostics) as the next two most important parameters, behind *α*. The top three parameters for Scenarios 3 and 4 are the same as for Scenario 2, for both PRCC and eFAST.

## Discussion

Accurate diagnostics can definitely guide targeted treatment. But for the diagnostic method to be effective in guiding decisions, preventing community impact and impacting situational awareness, several parameters other than sensitivity and specificity need to be considered. These include, but are not limited to, cost, speed to result, relative ease of use, and requirement of trained and experienced personnel. In this study, we have evaluated the impact of deployment of diagnostics for invasive NTS infection in a resource limited population and assessed a) impact on the individual and b) impact on the population, especially with respect to improved situational awareness and spread of antimicrobial resistance. In contrast to current understanding, full deployment of diagnostics resulting in targeted antibiotic use, resulted in a 50-90% reduction in total costs in our model, where the cost of diagnostics and antibiotics are also included (Figure 4). We inflated the cost of resistant antibiotic treatment to reflect the extreme un-desirability of multi-drug resistant strains, and to highlight the cost of resistant cases to both individuals and public health. We did not explicitly assign a cost to morbidity or mortality, but did quantify the change in number of deaths from NTS, and the size of an outbreak under considered scenarios. As has been observed for *Salmonella Typhi* (typhoid), our study shows that early detection and appropriate treatment are much more effective and cheaper than the status quo^9, 17, 42, 43^.

**Figure 4.**
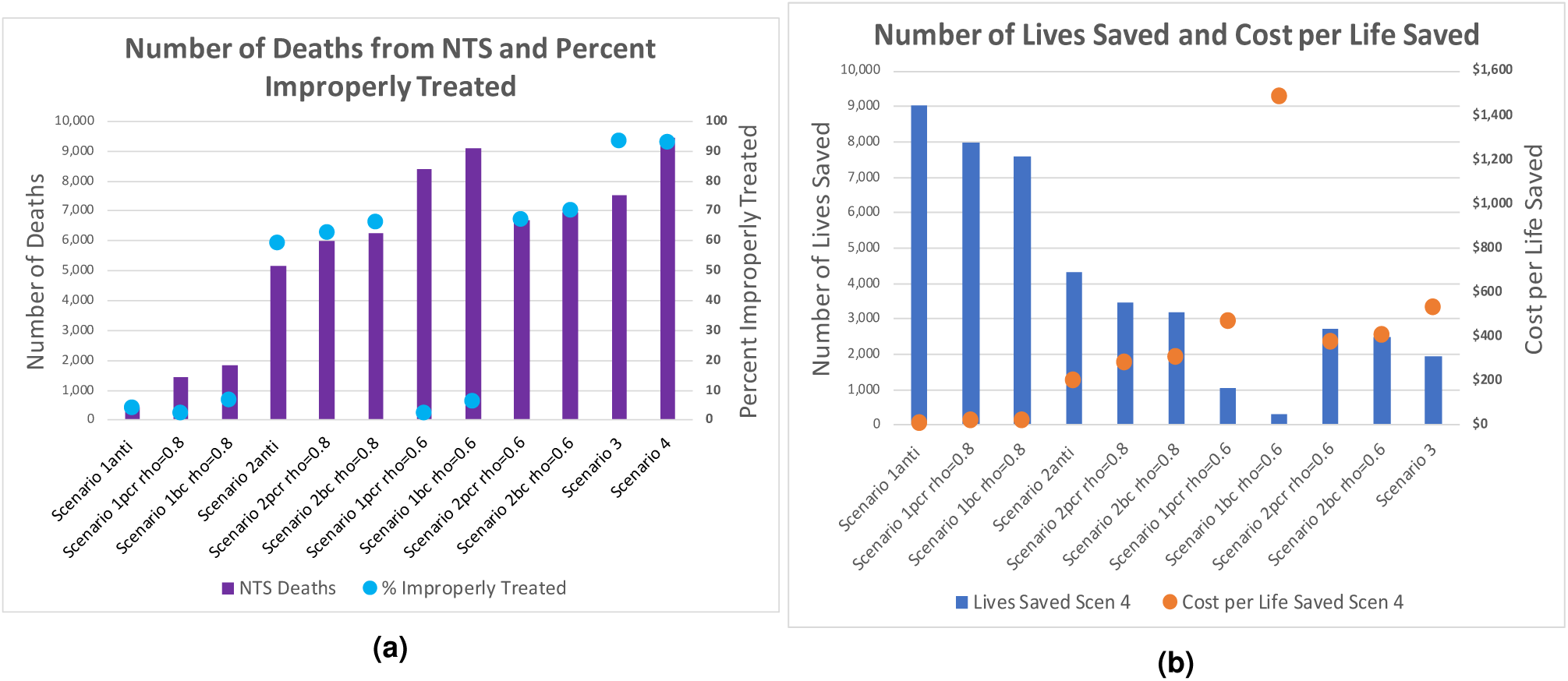
Subfigure 4a compares the number of deaths resulting from NTS with the percent of cases that are improperly treated. Number of deaths (left axis, purple bars) increases with percent of improper treatment (right axis, blue dots) and with time to POC result and patient compliance (“rho”). Subfigure 4b shows the number of lives saved (left axis, blue bars) in comparison to Scenario 4 and the cost per life saved (right axis, orange dots). The cost per life saved generally decreases with increased POC diagnostic deployment, targeted treatment, and patient compliance. If *ρ* = 0.6, or only 60% of patients return for diagnostic results and treatment the next day, then it is better to treat the severely symptomatic right away and reserve diagnostics for mild disease.

We found that deploying point of care diagnostics with resulting targeted antibiotic use almost always results in both lives saved, and a total reduction in cost, as well as a smaller community-wide outbreak. Specifically, our findings indicate that rapid POC diagnostics that can guide therapuetic intervention with relevant antimicrobials in a timely manner are associated with maximal benefit to both the patient and the community at large. There is evidence in Kenya that freely available health care and proper treatment can reduce the prevalence of antibiotic-resistant strains of NTS^16^. Many lives can be saved by even partial deployment of reasonably effective diagnostics (with moderate sensitivity and specificity). Once diagnostics are deployed and used regularly, further increases in sensitivity and specificity can result in gains in terms of lives saved. The highest percentage of improperly treated in full diagnostic deployment (Scenario 1) is still lower than the lowest percentage of improperly treated in partial diagnostic deployment (Scenario 2). In particular, the use of an antibody diagnostic in Scenario 1 resulted in the lowest percentage of improperly treated patients of any of the diagnostics in any scenario at 4.2%, as well as the lowest total cost of any Scenario at $65k. Further contributing to the advisability of Scenario 1 with the antibody diagnostic is the lowest number of deaths from NTS during the outbreak, at 429. However, if *ρ* = 0.6 so only 60% return for targeted treatment, then Scenario 2 where severe cases are treated right away and diagnostics are reserved for mild symptoms, is more advisable for BC and PCR.

Spread of antimicrobial resistance is minimized when proper treatment is administered and the corollary is true with improper or unnecessary use of antibiotics. Thus, high percentages of improperly treated patients are not desirable. Scenarios 3 and 4, with no diagnostics deployed, have high percentages of improperly treated people. The highest total cost of any of the scenarios with the most deaths from disease (see Table 1) is also evident in these cases. Even with partial diagnostic deployment (Scenario 2) if *ρ* = 0.8, lives are saved regardless of the diagnostic used (Table 1 and Figure 4), compared to no diagnostic deployment (Scenarios 3 and 4) (reduced by 26 - 54%). Full POC diagnostic deployment (Scenario 1) that can not only identify the cause, but also provide therapeutic intervention at the time of the visit, reduces the number of deaths significantly. Even with diagnostics that are more time consuming, full-deployment results in significantly lower death rate with a 45% reduction for a culture diagnostic, a 60% reduction for PCR, and a 96% reduction in deaths compared to Scenario 4 for antibody when *ρ* = 0.8. Total cost of treatment for BC and PCR in Scenario 2 is higher than total cost of treatment for Scenario 4, however (Table 2 and Figure 4). Counter-intuitive to expectations, our results show that the full deployment of diagnostics (Scenario 1, *ρ* = 0.8) is in fact, cheaper than all other scenarios, and is the only scenario where cost for properly treated outweighs cost for improperly treated (Table 2 and Figure 4). If *ρ* ≤ 0.6 then Scenario 2 is better than Scenario 1 with BC and PCR since less than 60% of people with iNTS will be treated under Scenario 1 (BC and PCR). It would be interesting in future work to consider a hybrid version of Scenarios 1 and 2 for BC and PCR.

For full and partial deployment of POC diagnostics, bacterial culture and PCR diagnostics were found to present with higher percentages of improper treatment than the antibody-based methods, suggesting that the the former two approaches do not adequately minimize antibiotic resistance. While some of this is due to lower sensitivity and specificity, our analysis suggests that the biggest factor is the time it takes to get a result back, which is directly related to *ρ*, the proportion of people who return to get diagnostic results and targeted treatment. Rapid diagnostics are most effective in our model to minimize impact and mitigate spread. Additionally, the percent of improperly treated NTS cases contributed significantly to higher total costs at both an individual and population level.

Apart from diagnostic use, the number of deaths from NTS can be reduced most significantly by decreasing the transmission rate (*α*). When full diagnostics are deployed (Scenario 1), increasing *κ*_*a*_, the rate at which the sensitive strain is cleared after receiving antibiotics, or decreasing the proportion of non-resistant NTS people who progress to invasive disease (*γ*) are the most effective ways to reduce deaths. In all other scenarios, it is more effective to increase 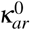, the rate at which the resistant strain is cleared after receiving broad spectrum and then resistant-appropriate antibiotics, with no diagnostics, or to decrease *β*, the proportion of NTS resistant individuals who are symptomatic. In the sensitivity analysis for Scenario 1 with antibody diagnostic (Table 3), *α, β*, *γ, κ*_*a*_, *λ*_*anti*_, and *σ*_*anti*_ have significant p-values in each of PRCC values and eFAST single index *S*_*i*_ and total index *S*_*ti*_ values. Once full diagnostics use with the antibody diagnostic (POC with very quick results, *ρ* = 1) is in place, the sensitive parameters indicate the next best areas for improvement. The current primary methods of reducing transmission are improved infrastructure and sanitation, education and surveillance, decreasing the carrier and reservoir populations, and use of a vaccination. However, these improvements that reduce transmission rate, *α*, take a long time to implement and are inherently tied to the socio-economic growth of the region, making a short-term solution necessary. The proportion of people who progress to invasive disease (*γ, β*) depends on the pathogen strain and on the health status of individuals in the community. Improving nutrition, decreasing malaria spread, reducing HIV prevalence, improving treatment for HIV and malaria, and generally improving public health access would reduce the number of invasive cases. However, as with transmission reduction, these are long-term goals that would take years to implement. Time to recovery after treatment, *κ*_*a*_, and the sensitivity and specificity, *λ*_*anti*_ and *σ*_*anti*_, can be improved with new drugs and better diagnostics. So, while the largest gain by far comes from implementing moderately accurate POC diagnostics and targeted treatment in the first place, improvements in diagnostic accuracy would improve outcomes even more. The rates at which a particular antibiotic clears invasive and non-invasive NTS, *κ*_*ar*_ and *κ*_*a*_ respectively, are among the most significant in each of our sensitivity analyses. So, as research in treatment progresses, and duration of antibiotic treatment needed to clear the pathogen decreases, the number of deaths from NTS will also decrease. Reducing this time, however, would require new drug discovery or approval of new dosages.

**Table 3.**
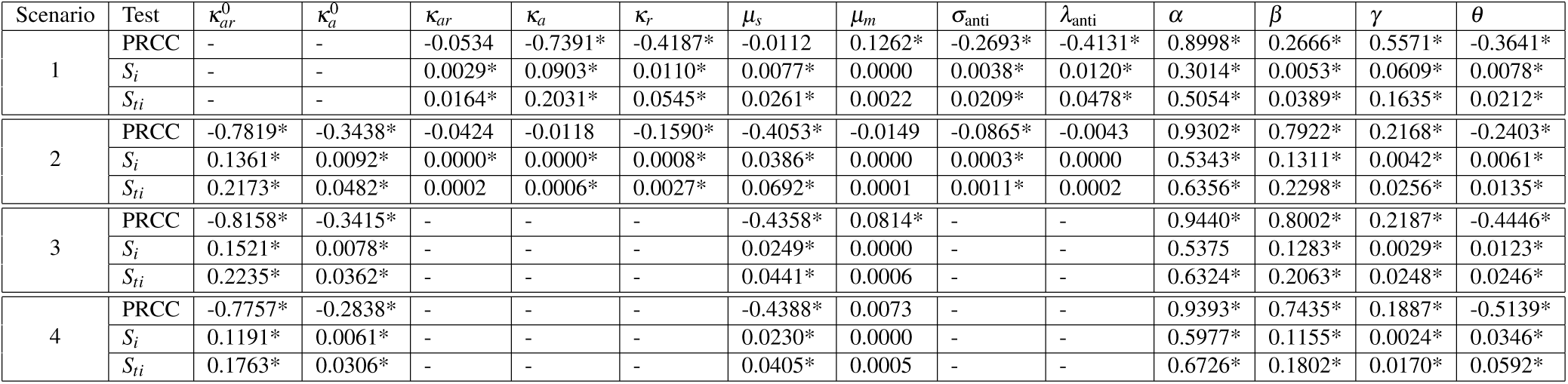
PRCC values, first- and total-order indices with their p-values for measuring the sensitivity of Scenario 1, 2, 3 and 4’s parameters to model R. Parameters were allowed to vary ±50% of their nominal values. The sample space was obtained using Latin Hypercube sampling. Values with a * have a p value less than 0.05. 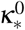 is the treatment/recovery rate when no diagnostic is used and *κ*_*_ is the rate with time to diagnostic result added (see Methods).

**Table 4.** Parameter descriptions, baseline values, parameter ranges, and references.

| Parameters | Definition | Baseline | Range |
| --- | --- | --- | --- |
| $\kappa_a$ | Broad Spectrum Antibiotic Treatment | $\frac{1}{14+\phi}$ | 14 days<sup>22</sup> |
| $\kappa_r$ | Resistant Antibiotic Treatment | $\frac{1}{14+\phi}$ | 10-14 days<sup>22</sup> |
| $\kappa_{ar}$ | Broad Spectrum then Resistant Treatment | $\frac{1}{28+\phi}$ | 24-28 days<sup>16,22</sup> |
| $\kappa_{ra}$ | Resistant Treatment then Broad Spectrum Treatment | $\frac{1}{28+\phi}$ | 24-28 days<sup>22</sup> |
| $\mu_s$ | Symptomatic Death Rate | 0.0195 | 0.0179-0.0446<sup>5,53</sup> |
| $\mu_m$ | Mildly Symptomatic Death Rate | 0.0038 | 0.00027 - 0.0073<sup>5,16,54</sup> |
| $\sigma_{PCR}$ | Diagnostic Sensitivity for PCR [0,1] | 0.95 | 90-100%<sup>10,23</sup> |
| $\lambda_{PCR}$ | Diagnostic Specificity for PCR [0,1] | 0.95 | 90-100%<sup>10,23</sup> |
| $\sigma_{BC}$ | Diagnostic Sensitivity for Bacteria Culture[0,1] | 0.79 | 79%<sup>10,23</sup> |
| $\lambda_{BC}$ | Diagnostic Specificity for Bacteria Culture [0,1] | 0.89 | 89%<sup>10,23</sup> |
| $\sigma_{AB}$ | Diagnostic Sensitivity for Antibody [0,1] | 0.89 | 78-100%<sup>10,23</sup> |
| $\lambda_{AB}$ | Diagnostic Specificity for Antibody [0,1] | 0.92 | 90-94%<sup>10,23</sup> |
| $\alpha$ | Infection Rate | 0.0000027 | $0.1*\alpha - 2*\alpha$<sup>5,16</sup> |
| $\beta$ | Proportion of Resistant NTS Severe[0,1] | 0.53 | 0.2-0.6<sup>2,16,55</sup> |
| $\theta$ | Natural Clearance Rate | 0.175 | 0.123 - 0.30<sup>5,16</sup> |
| $\gamma$ | Proportion of Non-resistant NTS Severe [0,1] | 0.47 | 0.2-0.5<sup>55</sup> |
| $\alpha_w$ | Infection Rate from Environment | 0 | $0.1\alpha - 2\alpha$<sup>5,16</sup> |
| $\psi$ | Environmental Source of NTS | 6 | [assumed] |
| $\tau$ | Shed Rate of Infectious into Environment | 0.01 | [assumed] |
| $\theta_w$ | Natural Clearance Rate of NTS in the Environment | 0.07 | 0.03–0.5<sup>55</sup> |
| $\rho$ | Fraction treated after receiving Antibody Diagnostic | 1.0 | 0.8-1.0 [assumed] |
|  | Fraction treated after receiving BC Diagnostic | 0.6 | 0.5-0.7 [assumed] |
|  | Fraction treated after receiving PCR Diagnostic | 0.6 | 0.5-0.7 [assumed] |

The transmission rate can be higher in small children and immuno-compromised adults and these rates vary even among these high-risk groups. We chose a baseline transmission risk determined from computations of incidence in the general population as a starting place to compute something like minimal risk for these groups. Our standard simplifying assumption of even mixing within the population generally results in higher overall transmission than is observed in the field, warranting conservative estimates of transmission. Our goal, then, was to calculate the impact of NTS and diagnostics/targeted treatment on a mixed population of high-risk people. In future models, it would be interesting to consider the high-risk groups separately to tease out the impact of targeted interventions in the sub-populations.

We also considered carriers or people with recrudescence and recurring infections that could act as sources for infecting the community in our model via an “Environmental” compartment. HIV-infected adults^35^ are at particular risk of becoming carriers. In fact, there is evidence that, unlike in the developed world, the NTS313 strain can be spread directly between humans rather than strictly *via* zoonoses, facilitated by carriers and immuno-depressed and immuno-naive individuals^44^. Maximum duration of shedding is believed to be 1 year for carriers and is 4-7 weeks for most other cases^14, 28^. The Environmental compartment also included zoonotic reservoirs, while the simplified model set transmission proportional to the number of people infected. As seen in Supplemental Material, the additional hosts served to increase the overall transmission, with magnitude depending on the density of additional reservoirs and carriers and their contact with susceptible high-risk hosts. It did not change general patterns in the effectiveness of diagnostic and treatment deployment. When reservoirs and carriers were not considered, our model did not exhibit long-term persistence of non-resistant NTS, but rather isolated outbreaks. This is consistent with observations in regions without significant carriers/reservoirs and with improved sanitation. However, it should be noted that in the absence of diagnostics, a 20% increase in the transmission rate, *α*, would bring the non-resistant strain *R*_0_ above 1 and that without the use of diagnostics, the resistant NTS strain has *R*_0_ > 1 and can persist long-term in the system without carriers. It would be interesting to consider carriers and reservoirs in more detail in order to understand the potential roles of antiretrovirals and sanitation on the system. However, further studies and data would be needed to parameterize such a detailed model.

NTS responds strongly to co-morbidities and risk factors such as HIV, malnutrition, malaria infection, and anemia, as well as rural settings and age^3, 5^. It would therefore be interesting to consider how addressing these co-morbidites and risk factors (i.e. treating malaria, antiretrovirals for HIV, improved nutrition) would change NTS dynamics. We only considered NTS in this study, but propose to extend this approach to include multiple circulating pathogens, including bacteria, viruses, and macro-parasites, that cause similar symptoms, to examine how a combination of empirical and diagnostic decision making could best be deployed in high-disease-burden areas in future studies. In Kenya, the most common diarrheal infections are caused by NTS, Rotaviruses, *Salmonella Typhi*, and *E*. *coli*^16^. There is currently no vaccine for NTS, but there has been research in this area, making including vaccination a possible extension to our model^45^. To further develop this model, we would like to explicitly quantify the cost in antibiotic resistance in some manner other than that represented by the cost of improper treatment. A potential way to do this would be to track the variability of percent of resistant cases across scenarios. There is also potential for the methods used here to be generalized and applied to other antibiotic-resistant bacteria case studies.

In the interest of clarity along with sparsity of relevant data, we made several assumptions in our model that may need to be relaxed or re-evaluated for a more accurate assessment for particular geography, populations, diagnostics, and available antimicrobials. We assumed that upon recovery, all at-risk patients returned to a fully susceptible state. In fact, there may be a period of immunity, which is supported by the fact that most healthy adults from endemic regions develop immunity from Salmonellosis. This assumption may result in model outbreaks moving faster than is observed in the field. However, in total numbers, our model output is comparable to an outbreak in Malawi^6^. We assumed everyone who was infected sought treatment. Since the actual rate of seeking treatment is confounded by several factors, including distance from health provider, access to health care, economic status, acquiring treatment from traditional healers or from non-licensed pharmaceutical vendors, among other, this assumption will need re-visiting based on more focused data from different populations. We began our simulations with equal numbers of drug-susceptible and drug-resistant infections and with a significant number of initial infections which may or may not be likely under normal conditions. This assumption also avoids the initial, highly stochastic, invasion of a strain. We assumed that, absent antibiotics, the resistant and sensitive strains are equally fit. Fitness of antimicrobial resistant strains is an evolving field of study, and current research suggests that while many pathogens acquire resistance to antimicrobials at a fitness cost, the human-adaptation and evolution of some strains may overcome these limitations^46, 47^. Finally, different culture, PCR, and antibody diagnostics have pros and cons that were not fully addressed in this study nor explicitly included in the parameter values. Some diagnostics can only detect certain strains, so must be targeted for the region considered, while others are more general and more likely to detect and emerging strain. Most likely, a combination of diagnostic methods that provide different information would be prudent in the field.

Despite these many assumptions, the core question of whether deployment of a diagnostic can improve patient outcomes and mitigate the spread of antibiotic resistance is effectively answered by our work. This is true because the assumptions impact each scenario equally. Each of these assumptions can be re-visited and investigated using our modeling approach to obtain more granularity on deployment of diagnostics in a chosen population. In conclusion, our analysis gives tantalizing evidence that POC diagnostic deployment coupled with improved treatment not only greatly reduces number of deaths and disease, but significantly mitigates overall costs associated with NTS. Our model shows that even imperfect diagnostics (e.g. under 95% specificity and sensitivity) are much better than none at all for the individual, and the community, and suggest that it is critical that the time to result and proper treatment should be minimized to improve outcomes. Diagnostics have the potential to provide important situational awareness for local, state, country and even global public health decision makers. Since no gold-standard diagnostics for NTS currently exist^10, 48^, we hope this research will motivate more investment in understanding NTS dynamics and developing point of care diagnostic capabilities.

## Methods

### Description of the Model

Systems of ordinary differential equations are used to model each of these scenarios by considering variations of a continuous-time compartmental SIR-type model. In this framework, patients are either Susceptible *(S*), Infectious with Resistant NTS (*I*), Infectious with Sensitive NTS (*J*), or Removed via death (*R*). To include other sources of NTS in the environment *W*_*I*_ and *W*_*J*_ are the environmental sources of resistant and sensitive NTS, respectively. The Susceptible compartment includes immuno-compromised (e.g. HIV positive) and immuno-naive (e.g. infants or young children) individuals that are at high risk for symptomatic NTS infections. The infectious compartment includes all infected and infectious individuals. The Removed via death compartment includes only individuals who have died as a result of NTS.

While the reality of immune dynamics is more complicated than our simplifying assumption, we found significant evidence that young children and HIV-positive people do not gain functional immunity to NTS. Even with previous exposure, children do not gain immunity until around 36 months of age^36, 49, 50^, particularly with common added factors of malaria, malnutrition, recent antimicrobial use, sickle cell, etc. HIV-positive people have also been observed to exhibit susceptibility to NTS after infection (recrudescence is also observed)^5, 51^. So, due to unknowns about permanent immunity in immuno-compromised or immuno-naive populations and the high number of strains circulating, individuals in the Infectious compartment who recover return to the Susceptible compartment^48^.

*S*(*t*), *I*_*_(*t*), *J*_*_(*t*), *R*(*t*) and *W*_*_(*t*) then give the population at time *t* that are in each of the compartments. The rate of movement between the compartments is dictated by a combination of relevant parameters which are described below^52^. In each scenario, patients are classified as either mildly symptomatic (subscript *m*, gastroenteritis) or severely symptomatic (subscript *s*, invasive with fever) with sensitive strains of NTS (denoted by *J*_*m*_ and *J*_*s*_) and resistant strains (denoted *I*_*m*_ and *I*_*s*_). ‘Sensitive’ strains of NTS respond to commonly prescribed broad-spectrum antimicrobials, while ‘resistant’ strains are resistant to one or more common antimicrobials.

The proportion of individuals infected with the resistant strain who are mildly symptomatic is represented by *β* and the proportion that are severely symptomatic is 1 − *β*. Likewise, the proportion of individuals infected with the sensitive strain who are mildly symptomatic is represented by *γ* and the proportion that are severely symptomatic is 1 − *γ*. The mildly symptomatic death rate for the high risk population, *µ*_*m*_, was calibrated to give a case fatality rate of 2% with a range of 0.15% −4%^54^. The symptomatic death rate, *µ*_*s*_, was calibrated to give a mean case fatality rate of about 22% for the non-resistant strain and 35% for the resistant in the high-risk population with no diagnostics used; a range of about 20% - 50% is observed in the literature^4, 5^.

We assumed that transmission of NTS in a community is proportional to the number of infectious individuals (*I*_*_, *J*_*_) at any given time. Additionally, we modelled environmental sources of NTS such as zoonotic reservoirs and human long-term carriers as the compartments *W*_*I*_ and *W*_*J*_. We calibrated the transmission rate, *α*, to correspond to an incidence of about 200 people per 100,000 per year^3^. Computed incidence varies significantly across studies and regions, probably due to a combination of limited data and variation in the high- and low-risk populations. Incidence in the general population ranges from 1.4 − 2, 520 per 100k^4^, was 175-388 per 100k in children in the same study and 1800-9000 per 100k in HIV positive adults^51^. Feasy et al. 2012 found similar numbers with 2000-7500 per 100k in HIV positive adults^5^. Mandomando et al. 2009 found iNTS incidence of 240/100k and 108/100k^56^. To capture this variation, we considered a wider range of transmission rates in our sensitivity analysis. We let the transmission rate, *α*, be the same for both resistant and non-resistant strains. The environmental transmission rate, *α*_*w*_, has been found to range anywhere from 0.1*α* to 2*α* in environmental transmission models^57–60^. We found no models for environmental transmission of NTS in humans. Though we analyzed our model outputs for this entire range, the qualitative nature of the model results did not change from the case when *α*_*w*_ = 0. Therefore, for simplicity, the results reported above correspond to the case when *α*_*w*_ = 0.

In our model, *κ*_*_ is the rate pathogen clearance from treatment, where 1*/κ*_*_ is the average time for course of treatment and full recovery. There are two types of treatment: a general antibiotic which successfully treats the sensitive strains of NTS, and is denoted by parameters with the subscript *a*, and the resistant treatment which treats resistant strains of NTS and is denoted by subscript *r*. When the strain of NTS is not responsive to the first treatment, the other treatment is applied and the treatment rates become *κ*_*ar*_, or the rate at which a patient incorrectly receives the standard sensitive strain antibiotic treatment then the proper resistant treatment, and *κ*_*ra*_, the rate at which a patient incorrectly receives the resistant treatment then the standard antibiotic treatment. In some cases, the resistant treatment will also clear sensitive strain, in which case *κ*_*ra*_ = *κ*_*r*_. A treatment procedure which takes a longer amount of time (e.g. use of a broad spectrum antibiotic and then a resistant treatment) would have a lesser *κ* value, and thus move individuals from the Infectious compartment to Susceptible (i.e. infection cleared) at a slower rate. We assume that everyone who needs treatment is given treatment. When diagnostics are used, we add the diagnostic result time to the total treatment time (Tables 5 and 4). For example, if it takes an average of 14 days for the individual to recover after being given standard antibiotic treatment and clinic receives the diagnostic result in 1 day, then the recovery rate, *κ*_*a*_, for that treatment would be 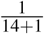. When diagnostics are not used, we will not add the diagnostic result time to the time to clearance; the treatment and clearance rates without diagnostics are referred to as 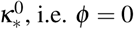, i.e. *ϕ* = 0. For the example above, 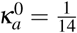.

**Table 5.** Time until diagnostic results received by clinic, assumed cost of each diagnostic, and assumed cost of antibiotic treatment. Costs or antibiotics are estimated based on averages and capture the relative cost of first-line antibiotics and antibiotics effective against resistant strains^16^.

| Parameter | Diagnostic | Time (days) | Cost (USD) |
| --- | --- | --- | --- |
| $\phi$ | Antibody | 0.0104 | 1 |
|  | BC | 1 | 5 |
|  | PCR | 1 | 10 |
| Broad Spectrum Antibiotic Treatment Course |  |  | 8.38 |
| Resistant Antibiotic Treatment Course |  |  | 62.50 |

The parameter *σ* corresponds to the sensitivity of the POC diagnostic; this is the proportion of resistant NTS infections that the diagnostic correctly determines, resulting in the patient receiving the proper treatment. The remaining (1 − *σ*) will initially receive the improper treatment and then receive proper treatment with rate constant *κ*_*ar*_. The parameter *λ* corresponds to the specificity of the diagnostic; this is the proportion of non-resistant NTS infections which the diagnostic determines. The remaining (1 − *λ*) will receive the improper treatment. All terms involving the use of a diagnostic before treatment are multiplied by a compliance constant, *ρ*. Because diagnostics are time-consuming, patients are at risk for leaving the clinic post-diagnostic and not returning and *ρ* accounts for the probability that a patient returns and seeks treatment given time to diagnostic result.

The natural clearance rate, *θ*, is the rate at which the body clears NTS without the help of an antibiotic. For our model, the severely symptomatic compartments (invasive disease) have a natural clearance rate of zero, meaning they require treatment to recover^5^. However mildly symptomatic people who do not die from the disease are assumed to recover without treatment, given time. 1*/θ* is the average time to natural recovery and pathogen clearance.

The shed rate *τ* is the rate of active shedding from infected individuals into the environment. An environmental source of NTS, *ψ* was used to include animal or human carriers. If the environment has approximately 1 million animals and humans, and 6% of the healthy animals and humans contract NTS and about 1% of those shed for several months, that equals about 600 individuals shedding. This value is then multiplied by the shedding rate *τ* to obtain a constant infusion. The clearance of NTS in the environment *θ*_*w*_ was titrated to achieve a steady state in both *W*_*I*_ and *W*_*J*_ that is greater than zero.

To maintain simplicity and keep results interpretable, the following assumptions are made in all four scenarios. Initial conditions (the starting populations) are denoted *S*(0), *I*_*s*_(0), *J*_*s*_(0), *I*_*m*_(0), *J*_*m*_(0),*W*_*I*_ (0),*W*_*J*_(0). Patients considered in this study are the young, elderly or immuno-compromised in a region similar to Kenya. Also, patients in any of the four Infectious compartments cannot move to other Infectious compartments; they are classified as either mildly symptomatic or severely symptomatic, and as either resistant or nonresistant, and cannot change classification during the course of their treatment plan. There are no long-term carrier classes, zoonotic or human, and no seasonality in the transmission rate. Rather than modeling long-term dynamics, we are considering a single outbreak in a closed population for about 3 years (1,000 days), which reflects observations in Africa^3^. We also assume that all people with NTS will report to a facility or health care provider that can perform a POC diagnostic and give a prescription for the appropriate treatment.

Below are the differential equations governing Scenario 1 with full diagnostic deployment and treatment for both symptomatic and mildly symptomatic:

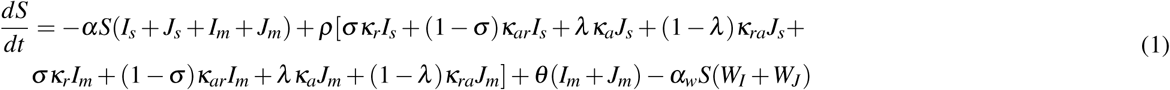

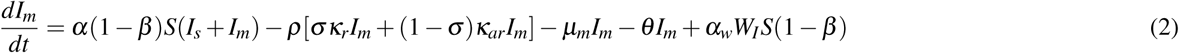

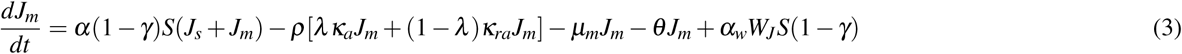

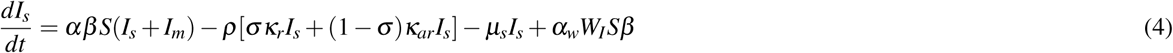

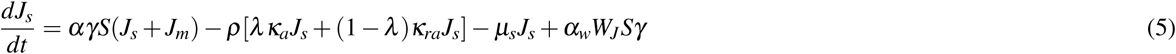

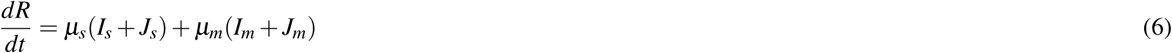

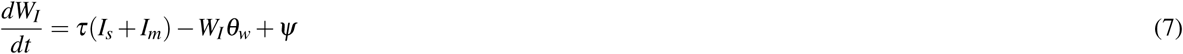

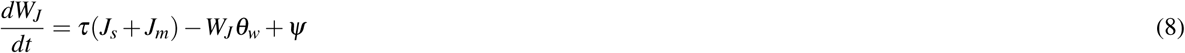

The equations for Scenarios 2 - 4 involve minor adaptations to Equations 1 - 8 and can be found in Supplementary Material. For the simplified model results presented in the main text, we set *α*_*w*_ = 0.

### Model Analysis and Simulations

#### Basic Reproduction Number

The basic reproduction number, *ℛ*_0_, for an infectious disease is the average number of secondary cases resulting from one infectious person introduced into a fully susceptible population. If *ℛ*_0_ > 1 then the pathogen can result in an outbreak, while if *ℛ*_0_ < 1 an outbreak is unlikely. It provides a measure for the rate at which a pathogen will spread in susceptible populations. We computed the basic reproduction number for our simplified model scenarios using the next generation method^61^. The **first term** of the basic reproduction numbers shown below is the *average number of secondary infections generated by a severely symptomatic individual* and the **second term** is the *average number of secondary infections generated by a mildly symptomatic individual*. Here, *N* is the total number of high-risk people in the population. For Scenario 1, the basic reproduction number for the non-resistant strain is

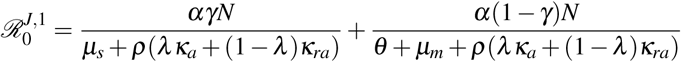

and for the resistant strain is

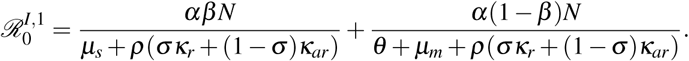

For Scenario 4, the basic reproduction number for the non-resistant strain is

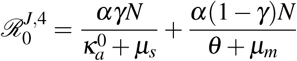

and for the resistant strain is

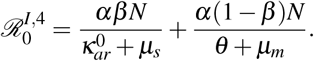

The remaining reproduction numbers are in the Supplementary Material. The basic reproduction numbers are directly proportional to transmission rate *α*. When diagnostics are deployed, the basic reproduction number is inversely proportional to the sensitivity (*σ*) and specificity (*λ*) of the diagnostics along with the compliance rate (*ρ*) and treatment rates (*κ*_*_) for resistant and non-resistant strains.

#### Simulations

We ran our model simulations for each of the scenarios at the baseline values found in Tables 5 and 4. We then assessed cost in terms of both total number of people who died from either invasive or non-invasive NTS and in terms of total dollars spent on diagnostics and treatment. We also assessed sensitivity of our model output to several parameter values. Scenarios 1–4 were simulated with each of the three possible diagnostics: PCR, BC, and antibody. Solutions were simulated using `ode45` in MATLAB and the following initial conditions:

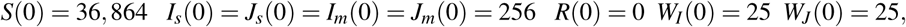

giving an initial high-risk population of 37,888 people, about 3.7% of a total population of 1,000,000 people. The differential equation solver ran for 1000 time units (days), producing results seen in Figures 1–2 and Tables 1–2.

#### Uncertainty Quantification

To find the sensitivity of the peak Death from NTS population (*D*) to changes in model input parameters, two different approaches were taken: Partial Rank Correlation Coefficients (PRCC) and extended Fourier Amplitude Sensitivity Testing (eFAST) (see^41, 62^). PRCC, a sampling based method, is the optimal analysis when nonlinear but monotonic relationships exist between inputs and outputs^62^, while eFAST, a variance based method, is used in models that utilize nonlinear and non-monotonic relationships between its input and output variables^63, 64^. PRCCs are a type of correlation coefficient that provide information regarding the amount of monotonicity that remains between the given input variable and the chosen output variable after linear effects of all but the chosen input variable have been removed. The sensitivity indices returned from eFAST, on the other hand, give a measure of the fractional variance that can be explained by individual input parameters or groups of parameters. First-order indices, *S*_*i*_, report the fraction of variance in the model output (peak value of *D* in our case) that can be explained by the variance in a given model input parameter. *S*_*Ti*_, the total-order index for a specific input parameter, reports the variance in model output (*R*) that remains when all variance caused by every other model input (every parameter *except* the given parameter) is removed.

PRCCs and eFAST sensitivity indices are included in Table 3 and Supplemental Material for Scenarios 1–4 respectively. For each of the model input parameters, values were generated between 50% above and below the values found in Table 4. Each of these generated values were drawn using a uniform distribution, and then PRCC and eFAST sensitivity indices were computed to determine sensitivity of the peak value of *R* to changes in these model input parameters. It is important to note that *ρ* was not included in the sensitivity analysis since it was an assumed quantity that we did not vary in this analysis.

We also performed an extended local sensitivity analysis on the basic reproduction number, *ℛ*_0_, to show how parameters affect the potential severity of an outbreak^65^. For this analysis, we examined the effect of changing *ρ*, as well as several other parameters, on the basic reproduction number. In particular, we set all other parameters to the baseline values and then varied the parameter being considered across its range. While this method of sensitivity analysis doesn’t capture the full variation that PRCC and eFAST analysis does, it allows for visualization of the role each parameter plays in the model as it varies. Results from this analysis are given in the Supplemental Material.

## Acknowledgements

CM was partially supported by NSF SEES Fellows grant CHE-1314029 and a LANL Director’s Postdoctoral Fellowship LDRD. We thank the PIC Math program. PIC Math is a program of the Mathematical Association of America (MAA) and the Society for Industrial and Applied Mathematics (SIAM). Support is provided by the National Science Foundation (NSF grant DMS-1345499). LANL is operated by Los Alamos National Security, LLC for the Department of Energy under contract DE-AC06NA25396. Approved for public release: LA-UR-. The content is solely the responsibility of the authors and does not necessarily represent the official views of the funding agencies. Thanks to Benjamin McMahon for many useful discussions and to D.J. Perkins for discussions and access to data.

## Author contributions statement

C.M., H.H., and H.M. conceived the study, C.M., H.H., T.G., A.C., and A.F. developed the model and model analysis methods, T.G., A.C., and A. F. wrote and ran the model simulation code, C.M., S.J., H.M. compiled parameter values from literature, H.M. and S.M. informed diagnostics, biology, and public health components of the model, C.M., T.G., S.J., H.M. compiled literature. C.M. wrote the manuscript draft. All authors reviewed the final manuscript.

## Supplemental Material

### Supplementary Material 1: Equations for Full Model

Note below that the *κ*^0^ notation means diagnostic time is NOT added to the total treatment time since diagnostics are not applied, i.e. *ϕ* = 0. Scenario 2 equations:

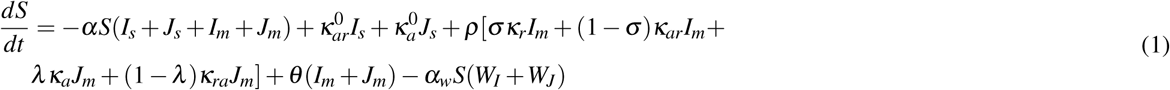

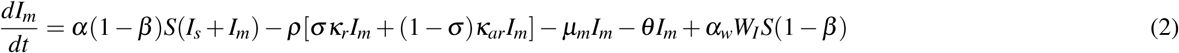

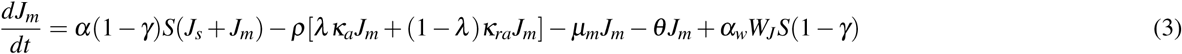

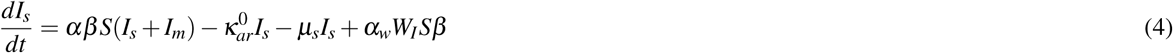

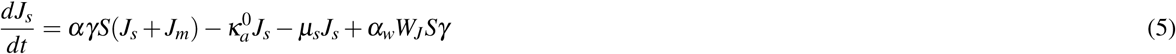

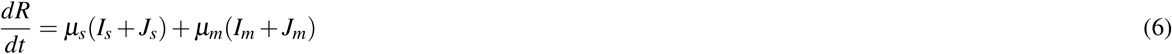

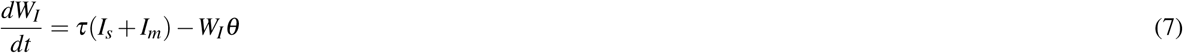

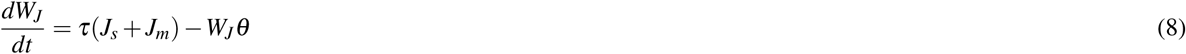

Scenario 3 equations:

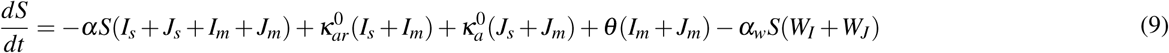

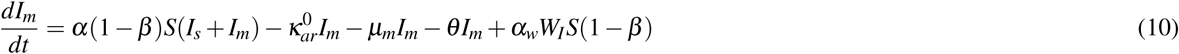

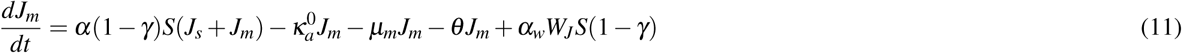

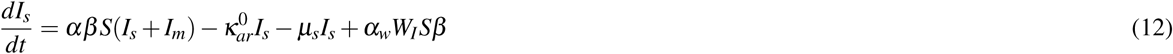

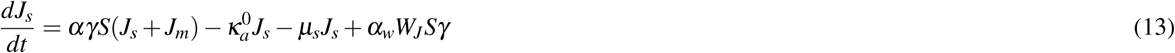

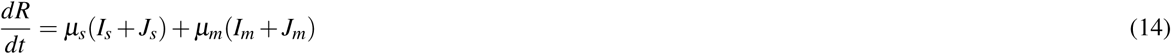

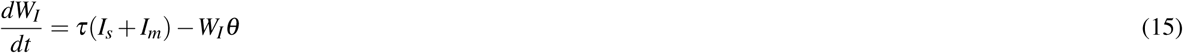

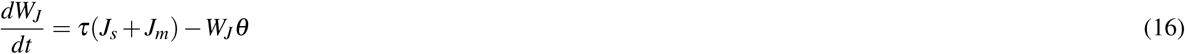

Scenario 4 equations:

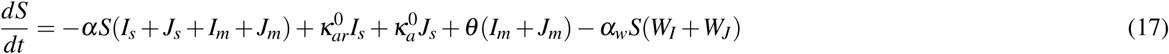

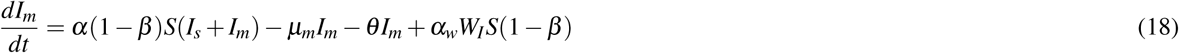

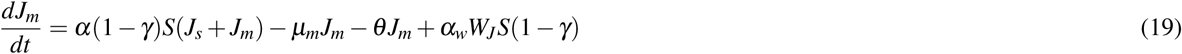

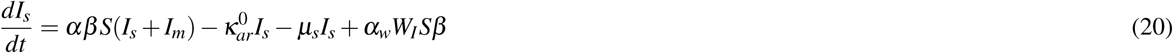

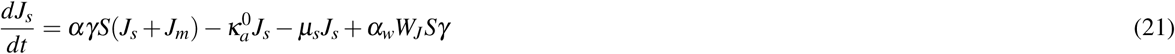

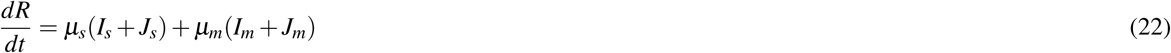

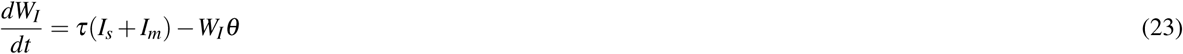

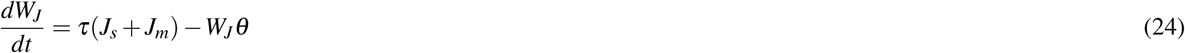

### Supplementary Material 2: Basic Reproduction Number for Simple Model (no Env compartment)

For Scenario 2, the basic reproduction number for the non-resistant strain is

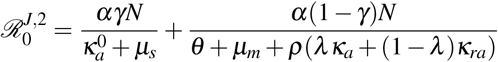

and for the resistant strain is

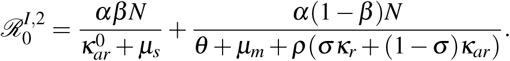

For Scenario 3, the basic reproduction number for the non-resistant strain is

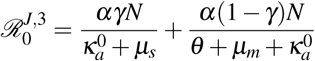

and for the resistant strain is

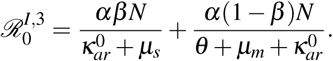

### Supplementary Material For Environmental/Outside Compartment Model

**Table 1.** Costs of diagnostic deployment and antibiotic use for each scenario with *α*_*w*_ = 0.5 * *α* where A is the cost of standard antibiotic treatment (effective on sensitive strain), R the cost of resistant strain treatment, and D the cost of the diagnostics. All costs are in U.S. Dollars (USD).

| Scenario | Diagnostic Used | Cost A (USD) | Cost AR (USD) | Cost DA (USD) | Cost DAR (USD) | Cost DR (USD) | Total Cost (USD) |
| --- | --- | --- | --- | --- | --- | --- | --- |
| 1 | Antibody | 0 | 0 | 115,448 | 59,618 | 422,811 | 597,877 |
|  | BC | 0 | 0 | 348,864 | 182,898 | 492,758 | 1,024,519 |
|  | PCR | 0 | 0 | 509,918 | 72,421 | 503,883 | 1,086,221 |
| 2 | Antibody | 44,339 | 1,114,363 | 15,007 | 30,434 | 647,765 | 1,851,908 |
|  | BC | 44,087 | 1,235,844 | 13,165 | 35,852 | 440,452 | 1,769,400 |
|  | PCR | 44,480 | 1,218,513 | 19,371 | 14,732 | 548,027 | 1,845,124 |
| 3 | None | 51,958 | 1,665,318 | 0 | 0 | 0 | 1,717,277 |
| 4 | None | 44,567 | 1,397,902 | 0 | 0 | 0 | 1,442,468 |

**Table 2.**
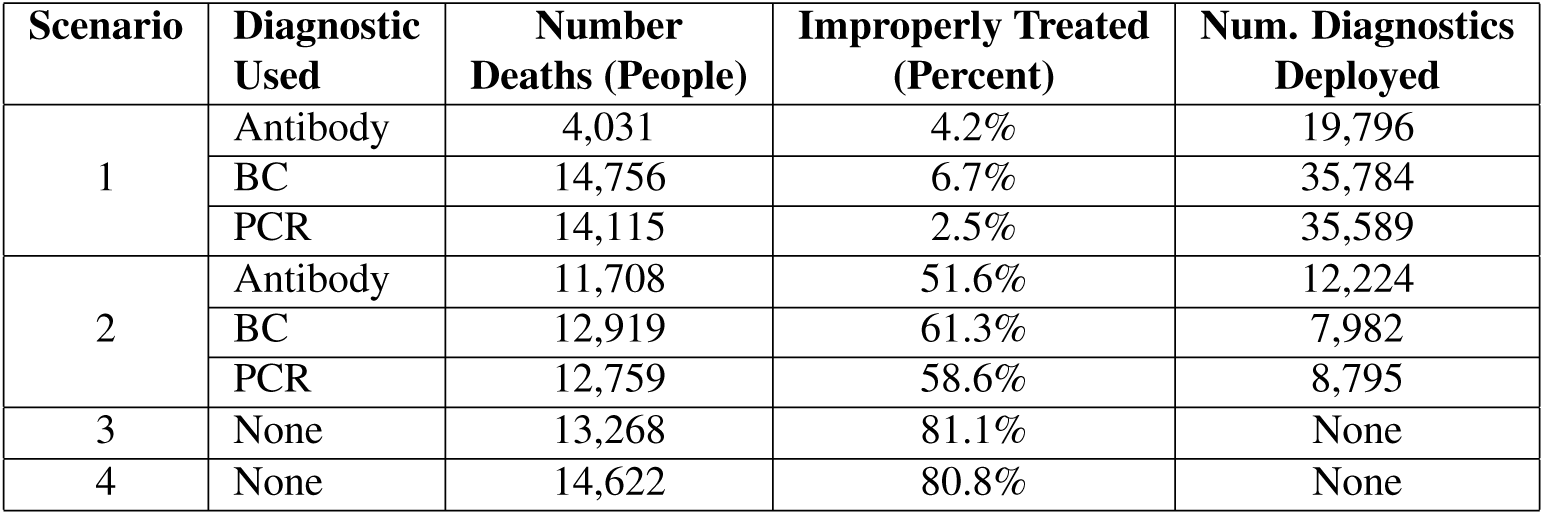
Number of deaths from NTS, percent of cases improperly treated, and the number of diagnostics used in each scenario with *α*_*w*_ = 0.5 \**α* run for 1,000 days. *ρ* = 0.6 for BC and PCR.

**Table 3.** PRCC values, first- and total-order indices with their p-values for measuring the sensitivity of Scenario 1, 2, 3 and 4’s non-environmental parameters to model R. Parameters were allowed to vary ± 50% of their nominal values. The sample space was obtained using Latin Hypercube sampling. Values with a * have a p value less than 0.05. Recall that 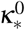 is the treatment/recovery rate when no diagnostic is used.

| Scenario | Test | $\kappa_{ar}^0$ | $\kappa_a^0$ | $\kappa_{ar}$ | $\kappa_a$ | $\kappa_r$ | $\mu_s$ | $\mu_m$ | $\sigma_{anti}$ | $\lambda_{anti}$ | $\alpha$ | $\beta$ | $\gamma$ | $\theta$ |
| --- | --- | --- | --- | --- | --- | --- | --- | --- | --- | --- | --- | --- | --- | --- |
| 1 | PRCC | - | - | -0.1309* | -0.7124* | -0.4163* | -0.0562 | 0.1102* | -0.2127* | -0.3795* | 0.8643* | 0.3319* | 0.6461* | -0.3615* |
| | $S_i$ | - | - | 0.0030* | 0.1003* | 0.0136* | 0.0018* | 0.0001 | 0.0039* | 0.0146* | 0.3093* | 0.0123* | 0.0811* | 0.0095* |
| | $S_{ti}$ | - | - | 0.0142* | 0.1642* | 0.0390* | 0.0122* | 0.0028 | 0.0166* | 0.0374* | 0.4026* | 0.0408* | 0.1417* | 0.0189* |
| 2 | PRCC | -0.7446* | -0.3471* | -0.0013 | 0.0252 | -0.0893* | -0.3551* | 0.0192 | -0.0472 | 0.0473 | 0.9078* | 0.8081* | 0.2898* | -0.3504 |
| | $S_i$ | 0.1445* | 0.0169* | 0.0000* | 0.0000* | 0.0008* | 0.0145* | 0.0000* | 0.0004* | 0.0000* | 0.4877* | 0.2000* | 0.0096* | 0.0075* |
| | $S_{ti}$ | 0.1902* | 0.0505* | 0.0002 | 0.0004* | 0.0021* | 0.0280* | 0.0001 | 0.0009 | 0.0002* | 0.5347* | 0.2589** | 0.0331* | 0.0123* |
| 3 | PRCC | -0.7856* | -0.3280* | - | - | - | -0.2978* | 0.0263 | - | - | 0.9188* | 0.7898* | 0.2269* | -0.4181* |
| | $S_i$ | 0.1527* | 0.0139* | - | - | - | 0.0133* | 0.0000* | - | - | 0.5098* | 0.1556* | 0.0068* | 0.0159* |
| | $S_{ti}$ | 0.1963* | 0.0405* | - | - | - | 0.0246* | 0.0002 | - | - | 0.5496* | 0.2004* | 0.0257* | 0.0229* |
| 4 | PRCC | -0.7550* | -0.3057* | - | - | - | -0.2755* | 0.0512 | - | - | 0.9216* | 0.7531* | 0.2531* | -0.4931* |
| | $S_i$ | 0.1303* | 0.0124* | - | - | - | 0.0113* | 0.0000* | - | - | 0.5430* | 0.1357* | 0.0055* | 0.0356* |
| | $S_{ti}$ | 0.1661* | 0.0364* | - | - | - | 0.0219* | 0.0002* | - | - | 0.5770* | 0.1746* | 0.0213* | 0.0484* |

**Table 4.**
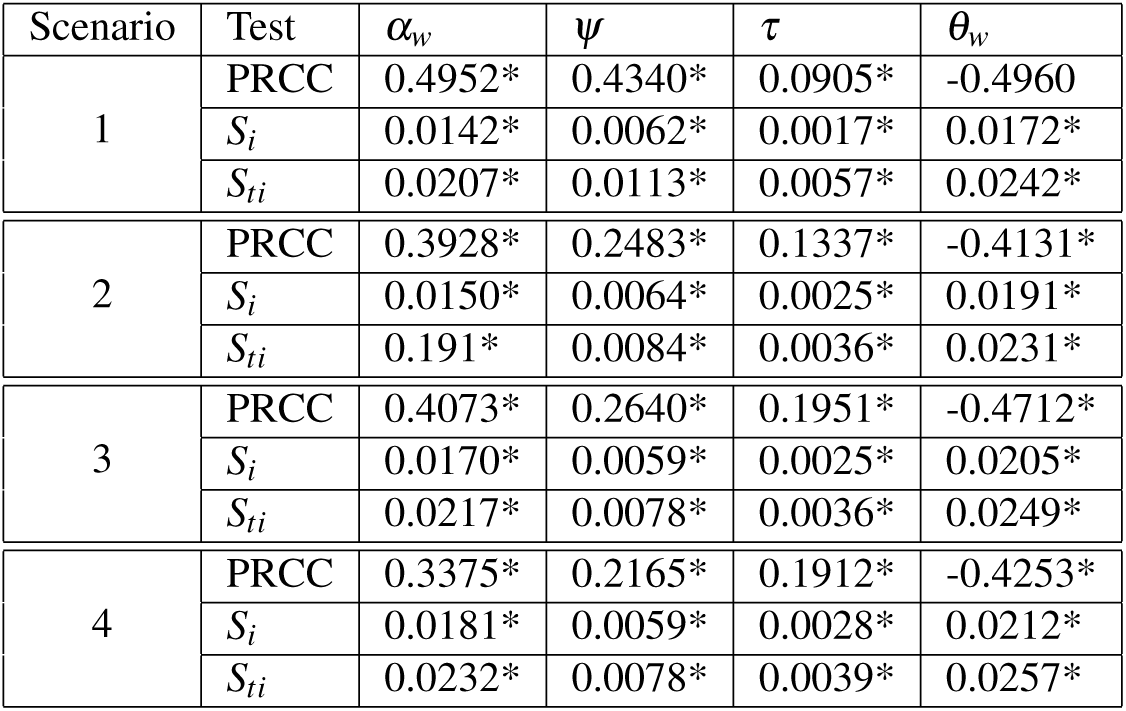
PRCC values, first- and total-order indices with their p-values for measuring the sensitivity of Scenario 1, 2, 3 and 4’s environmental parameters to model R. Parameters were allowed to vary ± 50% of their nominal values. The sample space was obtained using Latin Hypercube sampling. Values with a * have a p value less than 0.05.

**Figure 1.**
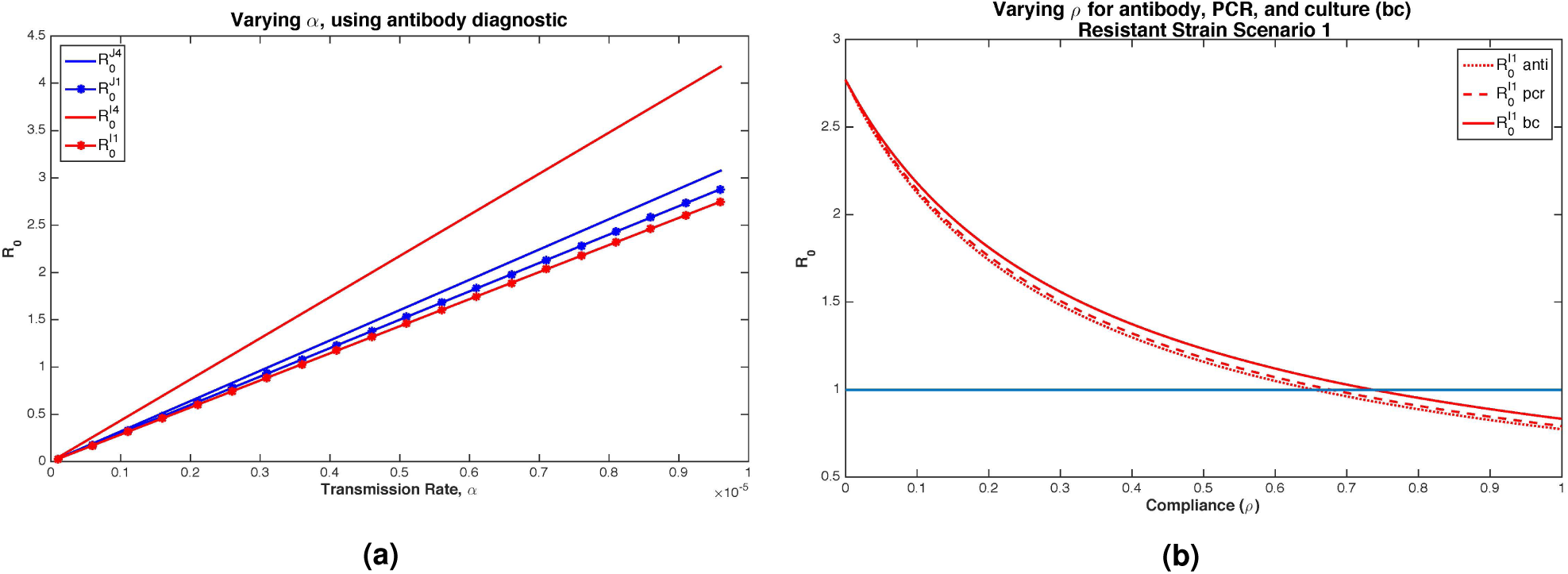
Sensitivity of *R*_0_ components to the transmission rate, *α* (Figure 1a), for different diagnostics and Scenarios 1 and 4, and to the compliance rate, *ρ* (Figure 1b). In subfigure 1a, blue lines are the sensitive strain and red the resistant strain. The flat lines are in the absence of diagnostics (Scenario 4) and starred lines with full diagnostic deployment (Scenario 1). In subfigure 1b, the compliance rate is the proportion of people who return to the clinic to receive diagnostic results and an appropriate treatment based on those results. *ℛ*_0_ increases linearly with the transmission rate and decreases non-linearly with compliance. The sensitivity of *ℛ*_0_ on compliance does not depend on diagnostic type.

**Figure 2.**
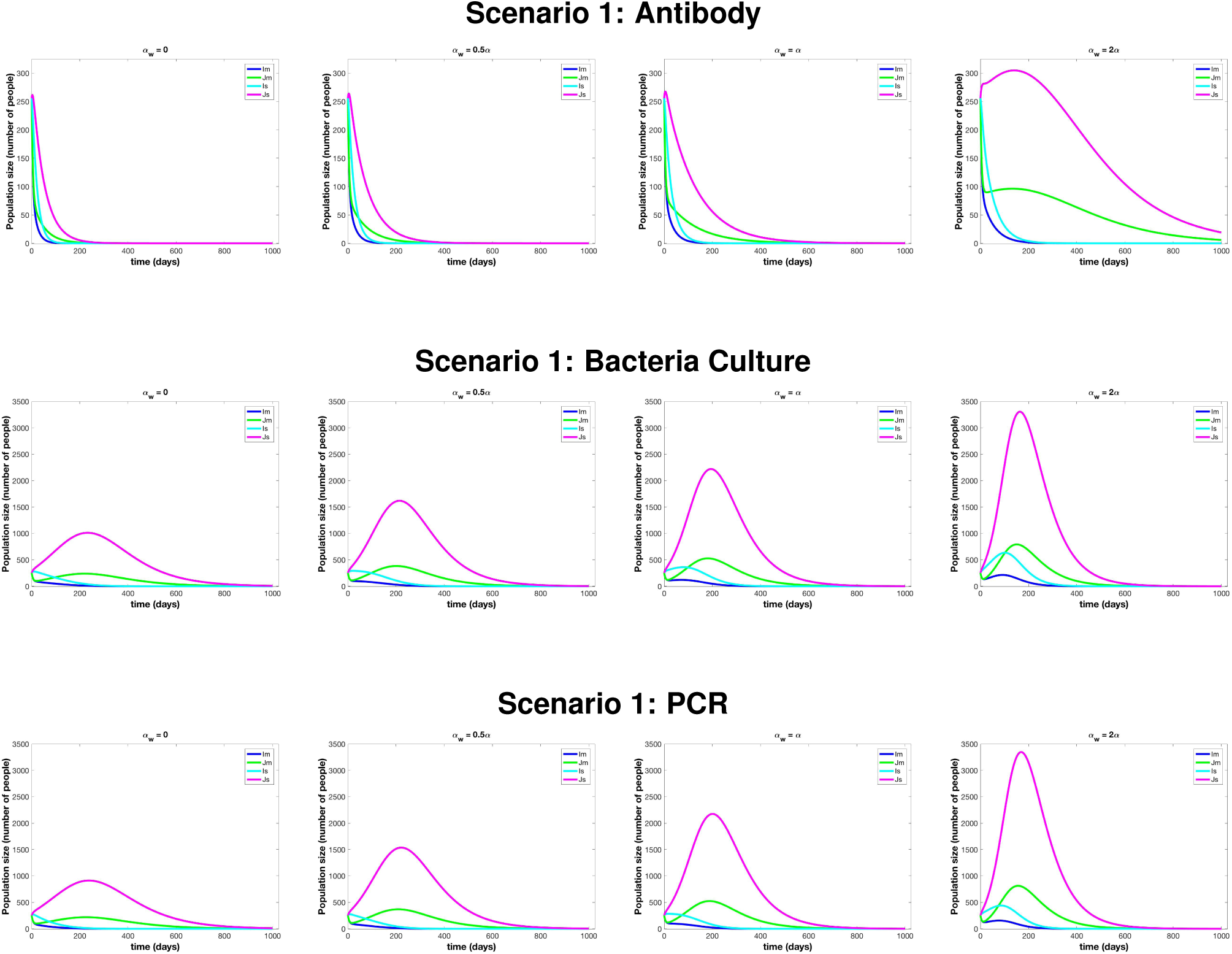
Outcome change with the environmental/low-risk compartment as the outside transmission rate, *α*_*w*_, changes for Scenario 1. While magnitude changes the general patterns remain the same except for very high values of outside transmission.

**Figure 3.**
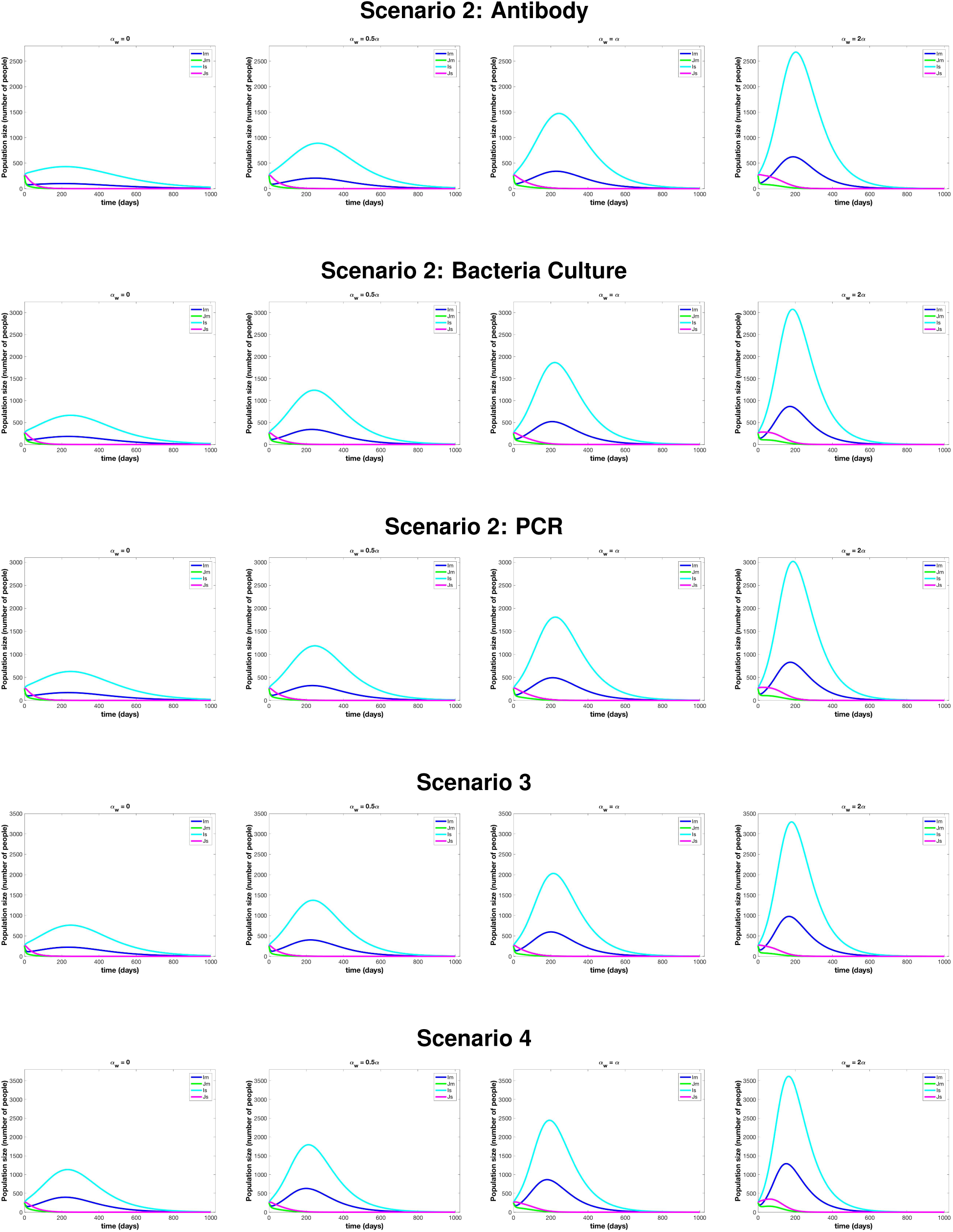
Outcome change with the environmental/low-risk compartment as the outside transmission rate, *α*_*w*_, changes for Scenarios 2 - 4. While magnitude changes the general patterns remain the same except for very high values of outside transmission.

**Figure 4.**
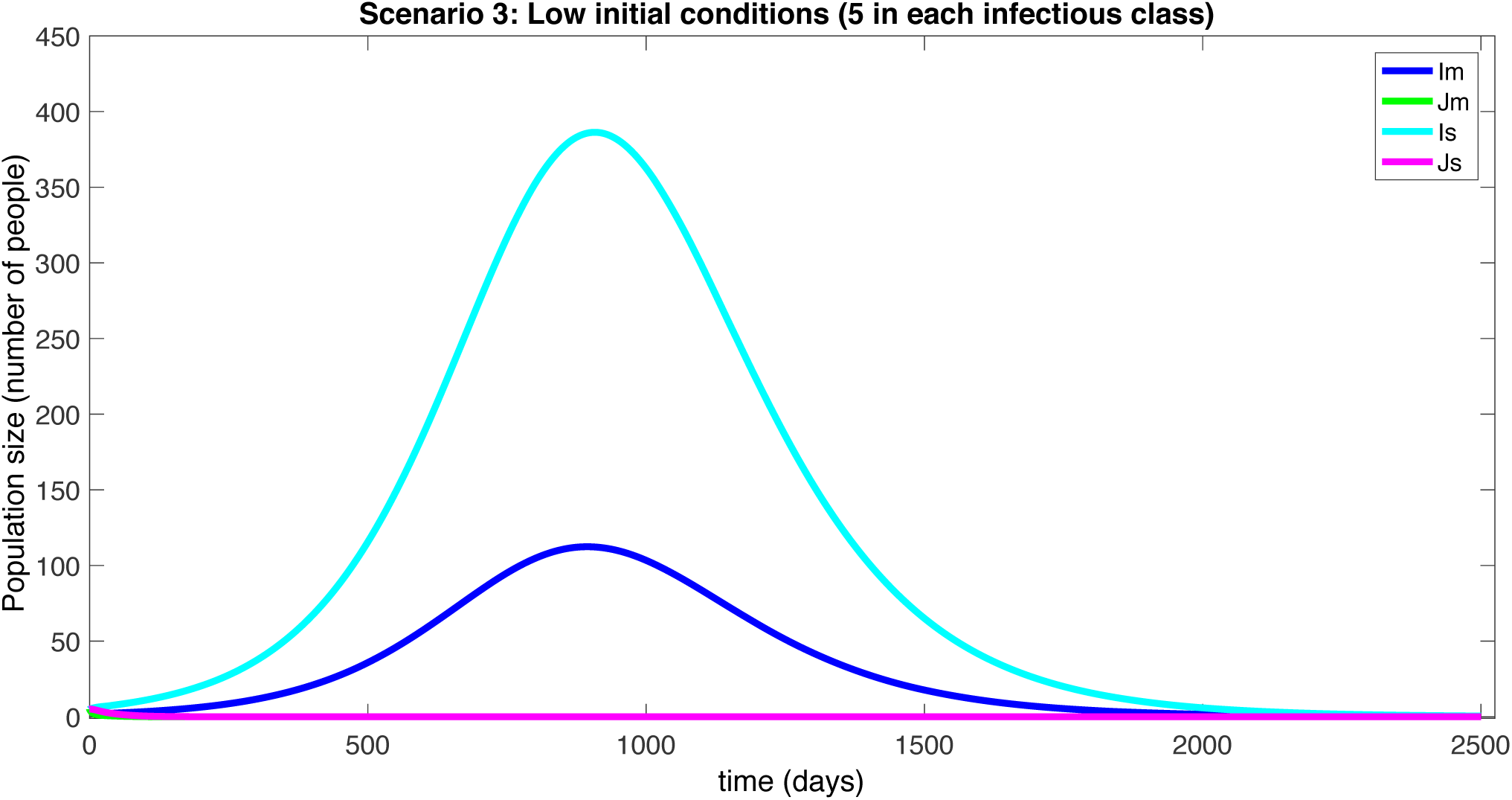
Scenario 3 for low initial conditions to simulate the outbreak in Blantyre, Malawi.

## References

1. Majowicz, S. E. et al. The global burden of nontyphoidal Salmonella gastroenteritis. Clin. Infect. Dis. 50, 882 (2010). DOI 10.1086/650733. /oup/backfile/Content_public/Journal/cid/50/6/10.1086/650733/2/50-6-882.pdf.

2. Lokken, K. L., Walker, G. T. & Tsolis, R. M. Disseminated infections with antibiotic-resistant non-typhoidal Salmonella strains: contributions of host and pathogen factors. Pathog. Dis. 74, 103 (2016).

3. Marks, F. et al. Incidence of invasive Salmonella disease in sub-saharan africa: a multicentre population-based surveillance study. The Lancet Glob. Heal. 5, e310–e323 (2017).

4. Uche, I. V., MacLennan, C. A. & Saul, A. A systematic review of the incidence, risk factors and case fatality rates of invasive nontyphoidal Salmonella (iNTS) disease in Africa (1966 to 2014). PLOS Neglected Trop. Dis. 11, 118 (2017).

5. Feasey, N. A., Dougan, G., Kingsley, R. A., Heyderman, R. S. & Gordon, M. A. Invasive non-typhoidal Salmonella disease: an emerging and neglected tropical disease in Africa. The Lancet 379, 2489–2499 (2012). DOI http://dx.doi.org/10.1016/S0140-6736(11)61752-2.

6. Gordon, M. A. et al. Epidemics of invasive salmonella enterica serovar enteritidis and s. enterica serovar typhimurium infection associated with multidrug resistance among adults and children in malawi. Clin. Infect. Dis. 46, 963–969 (2008).

7. Okoro, C. K. et al. Intracontinental spread of human invasive salmonella typhimurium pathovariants in sub-saharan africa. Nat. genetics 44, 1215–1221 (2012).

8. Were, T. et al. Bacteremia in kenyan children presenting with malaria. J. clinical microbiology 49, 671–676 (2011).

9. Oneko, M. et al. Emergence of community-acquired, multidrug-resistant invasive nontyphoidal salmonella disease in rural western kenya, 2009-2013. Clin. Infect. Dis. 61, S310–S316 (2015).

10. Andrews, J. R. & Ryan, E. T. Diagnostics for invasive salmonella infections: Current challenges and future directions. Vaccine 33, C8–C15 (2015). Global Progress on use of Vaccines for Invasive Salmonella Infections.

11. Reddy, E. A., Shaw, A. V. & Crump, J. A. Community-acquired bloodstream infections in africa: a systematic review and meta-analysis. The Lancet Infect. Dis. 10, 417–432 (2010).

12. Nadjm, B. et al. WHO guidelines for antimicrobial treatment in children admitted to hospital in an area of intense Plasmodium falciparum transmission: prospective study. BMJ 340 (2010).

13. Mabey, D., Peeling, R. W., Ustianowski, A. & Perkins, M. D. Diagnostics for the developing world. Nat. Rev. Microbiol. 2, 231–40 (2004). Copyright - Copyright Nature Publishing Group Mar 2004; Last updated - 2014-04-21.

14. Andoh, L. A. et al. Prevalence and characterization of Salmonella among humans in Ghana. Trop. Medicine Heal. 45, 3 (2017). DOI 10.1186/s41182-017-0043-z.

15. Lunguya, O. et al. Antimicrobial resistance in invasive non-typhoid Salmonella from the Democratic Republic of the Congo: emergence of decreased fluoroquinolone susceptibility and extended-spectrum beta lactamases. PLOS Neglected Trop. Dis. 7, e2103 (2013).

16. Group, K. W. Situation analysis and recommendations: Antibiotic use and resistance in Kenya (2017). URL https://www.cddep.org/sites/default/files/garp/sitan/pdf/garp-kenya.pdf.

17. Watson, C. H. & Edmunds, W. J. A review of typhoid fever transmission dynamic models and economic evaluations of vaccination. Vaccine 33, C42–C54 (2015).

18. Bakach, I., Just, M. R., Gambhir, M. & Fung, I. C.-H. Typhoid transmission: a historical perspective on mathematical model development. Transactions The Royal Soc. Trop. Medicine Hyg. 75 (2015).

19. Cvjetanović, B., Grab, B. & Uemura, K. Epidemiological model of typhoid fever and its use in the planning and evaluation of antityphoid immunization and sanitation programmes. Bull. World Heal. Organ. 45, 53 (1971).

20. Feasey, N. A. et al. Modelling the contributions of malaria, HIV, malnutrition and rainfall to the decline in paediatric invasive non-typhoidal Salmonella disease in Malawi. PLOS Neglected Trop. Dis. 9, 1–12 (2015). DOI 10.1371/journal.pntd.0003979.

21. Zhou, L. & Pollard, A. J. A fast and highly sensitive blood culture PCR method for clinical detection of Salmonella enterica serovar typhi. Annals Clin. Microbiol. Antimicrob. 9, 14 (2010).

22. Crump, J. A., Sjölund-Karlsson, M., Gordon, M. A. & Parry, C. M. Epidemiology, clinical presentation, laboratory diagnosis, antimicrobial resistance, and antimicrobial management of invasive Salmonella infections. Clin. Microbiol. Rev. 28, 901–937 (2015).

23. Ley, B. et al. Assessment and comparative analysis of a rapid diagnostic test (tubex R) for the diagnosis of typhoid fever among hospitalized children in rural Tanzania. BMC Infect. Dis. 11, 147 (2011).

24. Keddy, K. H. et al. Sensitivity and specificity of typhoid fever rapid antibody tests for laboratory diagnosis at two sub-Saharan African sites. Bull. World Heal. Organ. 89, 640–647 (2011).

25. MacLennan, C. A. et al. The neglected role of antibody in protection against bacteremia caused by nontyphoidal strains of Salmonella in African children. The J. Clin. Investig. 118, 1553–1562 (2008).

26. Kenya Ministry of Health. Policy guidelines for management of diarrhoea in children below five years in Kenya (2017). URL http://guidelines.health.go.ke:8000/media/Policy_Guidelines_for_Management_of_Diarrhoea_in_Children_Below.pdf.

27. Organization, W. H. Who recommendations on the management of diarrhoea and pneumonia in HIV-infected infants and children: Integrated management of childhood illness (IMCI) (2017). URL http://apps.who.int/iris/bitstream/10665/44471/1/9789241548083_eng.pdf.

28. Acheson, D. & Hohmann, E. L. Nontyphoidal salmonellosis. Clin. Infect. Dis. 32, 263 (2001). DOI 10.1086/318457.

29. Sánchez-Vargas, F. M., Abu-El-Haija, M. A. & Gómez-Duarte, O. G. Salmonella infections: an update on epidemiology, management, and prevention. Travel. Medicine Infect. Dis. 9, 263–277 (2011).

30. Abdulraheem, I., Adegboye, A. & Fatiregun, A. Self-medication with antibiotics: Empirical evidence from a Nigerian rural population. Br. J. Pharm. Res. 11 (2016).

31. Macro, O. et al. Knbs and icf macro: Kenya demographic and health survey 2008–09 (2010).

32. Okeke, I. N. & Ojo, K. K. Antimicrobial Resistance in Developing Countries (Springer, 2010).

33. Afriyie, E. O. et al. Antibiotics availability and usage in health facilities: A case of the offinso-south municipality of Ghana (2015).

34. Im, J. et al. Prevalence of salmonella excretion in stool: a community survey in 2 sites, guinea-bissau and senegal. Clin. infectious diseases 62, S50–S55 (2016).

35. Gordon, M. A. et al. Non-typhoidal Salmonella bacteraemia among HIV-infected Malawian adults: high mortality and frequent recrudescence. AIDS 16, 1633–1641 (2002).

36. Morpeth, S. C., Ramadhani, H. O. & Crump, J. A. Invasive non-typhi salmonella disease in africa. Clin. Infect. Dis. 49, 606–611 (2009).

37. Kariuki, S., Gordon, M. A., Feasey, N. & Parry, C. M. Antimicrobial resistance and management of invasive salmonella disease. Vaccine 33, C21–C29 (2015).

38. Kariuki, S. et al. Lack of clonal relationship between non-typhi salmonella strain types from humans and those isolated from animals living in close contact. FEMS Immunol. & Med. Microbiol. 33, 165–171 (2002).

39. Dione, M. M. et al. Clonal differences between non-typhoidal salmonella (nts) recovered from children and animals living in close contact in the gambia. PLOS Neglected Trop. Dis. 5, e1148 (2011).

40. Panzner, U. et al. Utilization of healthcare in the typhoid fever surveillance in africa program. Clin. Infect. Dis. 62, S56–S68 (2016).

41. Marino, S., Hogue, I. B., Ray, C. J. & Kirschner, D. E. A methodology for performing global uncertainty and sensitivity analysis in systems biology. J. Theor. Biol. 254, 178–196 (2008).

42. Bahl, R. et al. Costs of illness due to typhoid fever in an indian urban slum community: implications for vaccination policy. J. Heal. Popul. Nutr. 304–310 (2004).

43. Poulos, C. et al. A cost-benefit analysis of typhoid fever immunization programmes in an indian urban slum community. J. Heal. Popul. Nutr. 311–321 (2004).

44. Kariuki, S. et al. Invasive multidrug-resistant non-typhoidal Salmonella infections in africa: zoonotic or anthroponotic transmission? J. Med. Microbiol. 55, 585–591 (2006).

45. Ferreira, R. B. et al. A highly effective component vaccine against nontyphoidal Salmonella enterica infections. mBio 6, 14–15 (2015).

46. Maharjan, R. & Ferenci, T. The fitness costs and benefits of antibiotic resistance in drug-free microenvironments encountered in the human body. Environ. Microbiol. Reports 9, 635–641 (2017).

47. Zhang, C.-Z. et al. Resistance mechanisms and fitness of salmonella typhimurium and salmonella enteritidis mutants evolved under selection with ciprofloxacin in vitro. Sci. Reports 7, 9113 (2017).

48. Arya, G. et al. Epidemiology, pathogenesis, genoserotyping, antimicrobial resistance, and prevention and control of non-typhoidal Salmonella serovars. Curr. Clin. Microbiol. Reports 4, 43–53 (2017).

49. Nyirenda, T. S., Mandala, W. L., Gordon, M. A. & Mastroeni, P. Immunological bases of increased susceptibility to invasive nontyphoidal salmonella infection in children with malaria and anaemia. Microbes infection (2017).

50. Nyirenda, T. S. et al. Sequential acquisition of t cells and antibodies to nontyphoidal salmonella in malawian children. The J. infectious diseases 210, 56–64 (2014).

51. Gordon, M. A. Invasive non-typhoidal salmonella disease–epidemiology, pathogenesis and diagnosis. Curr. Opin. Infect. Dis. 24, 484 (2011).

52. Just, W. & Callender, H. Differential equation models of disease transmission (2015). URL http://www.ohio.edu/people/just/IONTW/ModuleDE.pdf.

53. Dhanoa, A. & Fatt, Q. K. Non-typhoidal Salmonella bacteraemia: epidemiology, clinical characteristics and its’ association with severe immunosuppression. Annals Clin. Microbiol. Antimicrob. 8, 15 (2009).

54. Kosek, M., Bern, C. & Guerrant, R. L. The global burden of diarrhoeal disease, as estimated from studies published between 1992 and 2000. Bull. World Heal. Organ. 81, 197–204 (2003).

55. Kennedy, M. et al. Hospitalizations and deaths due to Salmonella infections, FoodNet, 1996–1999. Clin. Infect. Dis. 38, S142–S148 (2004).

56. Mandomando, I. et al. Invasive non-typhoidal salmonella in mozambican children. Trop. Medicine & Int. Heal. 14, 1467–1474 (2009).

57. Xiao, Y., Bowers, R. G., Clancy, D. & French, N. P. Understanding the dynamics of salmonella infections in dairy herds: a modelling approach. J. Theor. Biol. 233, 159–175 (2005).

58. Chapagain, P. et al. A mathematical model of the dynamics of salmonella cerro infection in a us dairy herd. Epidemiol. & Infect. 136, 263–272 (2008).

59. Eisenberg, M. C., Robertson, S. L. & Tien, J. H. Identifiability and estimation of multiple transmission pathways in cholera and waterborne disease. J. Theor. Biol. 324, 84–102 (2013).

60. Tien, J. H. & Earn, D. J. Multiple transmission pathways and disease dynamics in a waterborne pathogen model. Bull. Math. Biol. 72, 1506–1533 (2010).

61. Van den Driessche, P. & Watmough, J. Reproduction numbers and sub-threshold endemic equilibria for compartmental models of disease transmission. Math. Biosci. 180, 29–48 (2002).

62. Saltelli, A. et al. Global Sensitivity Analysis: the Primer (John Wiley & Sons, 2008).

63. Ratto, M., Pagano, A. & Young, P. State dependent parameter metamodelling and sensitivity analysis. Comput. Phys. Commun. 177, 863–876 (2007).

64. Tarantola, S., Gatelli, D. & Mara, T. A. Random balance designs for the estimation of first order global sensitivity indices. Reliab. Eng. & Syst. Saf. 91, 717–727 (2006).

65. Manore, C. A., Hickmann, K. S., Xu, S., Wearing, H. J. & Hyman, J. M. Comparing dengue and chikungunya emergence and endemic transmission in a. aegypti and a. albopictus. J. Theor. Biol. 356, 174–191 (2014).

